# Optimal routing to cerebellum-like structures

**DOI:** 10.1101/2022.02.10.480014

**Authors:** Samuel Muscinelli, Mark Wagner, Ashok Litwin-Kumar

## Abstract

The vast expansion from mossy fibers to cerebellar granule cells produces a neural representation that supports functions including associative and internal model learning. This motif is shared by other cerebellum-like structures, including the insect mushroom body, electrosensory lobe of electric fish, and mammalian dorsal cochlear nucleus, and has inspired numerous theoretical models of its functional role. Less attention has been paid to structures immediately presynaptic to granule cell layers, whose architecture can be described as a “bottleneck” and whose functional role is not understood. We therefore develop a general theory of cerebellum-like structures in conjunction with their afferent pathways. This theory predicts the role of the pontine relay to cerebellar cortex and the glomerular organization of the insect antennal lobe. It also reconciles theories of nonlinear mixing with recent observations of correlated granule cell activity. More generally, it shows that structured compression followed by random expansion is an efficient architecture for flexible computation.

## Introduction

In the cerebral cortex, multiple densely connected, recurrent networks process input to form sensory representations. Theoretical models and studies of artificial neural networks have shown that such architectures are capable of extracting features from structured input spaces relevant for the production of complex behaviors [1]. However, the vertebrate cerebellum and cerebellum-like structures including the insect mushroom body, the electrosensory lobe of the electric fish, and the mammalian dorsal cochlear nucleus, operate on very different architectural principles [2]. In these areas, sensory and motor inputs are routed in a largely feedforward manner to a sparsely connected granule cell layer, whose neurons lack lateral recurrent interactions. These features suggest that such areas exploit a different strategy than the cerebral cortex to form their neural representations, despite being involved in many adaptive behaviors.

Many theories have focused on the computational role of the expanded granule cell representation in the cerebellum and cerebellum-like systems [3–7]. However, these theories have assumed a set of independent inputs, neglecting the upstream areas that construct them. As we show, this assumption severely underestimates the learning performance of such systems for structured inputs. We hypothesized that limitations due to input correlations are overcome by the specialized regions presynaptic to granule cell layers that process inputs to facilitate downstream learning. These regions have an architecture that can be described as a “bottleneck.” In the mammalian cerebellum, inputs to granule cells originating from the cerebral cortex arrive primarily via the pontine nuclei in the brainstem, which compresses the cortical representation [8]. In the insect olfactory system, about 50 classes of olfactory projection neurons in the antennal lobe route input from thousands of olfactory sensory neurons to the roughly 2000 Kenyon cells in the mushroom body, the analogs of cerebellar granule cells. Other cerebellum-like structures exhibit a similar architecture [2].

Theoretical analyses have provided explanations for both the large expansion in the number of granule cells and their small number of incoming connections (in-degree) without explicitly modeling the areas upstream of granule cells [6,7]. On the other hand, some of these upstream areas have been studied in isolation from the downstream granule cell layer. A number of studies have focused on the function of the insect antennal lobe, as well as the olfactory bulb, an analogous structure in mammals. Some have proposed that its main function is to de-noise olfactory sensory neuron signals [9], while others have argued for whitening the statistics of these responses [10,11]. The pontine nuclei upstream of cerebellar granule cells have received less attention. Recent experiments suggest that the pontine nuclei not only relay the cortical representation received from layer-5 pyramidal cells but also integrate and reshape it [12].

Here, we use a combination of simulations, analytical calculations, and data analysis to develop a general theory of cerebellum-like structures and their afferent pathways. We propose that the bottleneck architecture of regions presynaptic to granule-like layers can be understood from the twofold goal of increasing dimensionality and minimizing noise. Our theory predicts that incoming weights to these regions should be tuned to the input statistics, and we show that this is particularly beneficial when input neurons are noisy and correlated. When applied to the insect olfactory system, our theory explains its glomerular organization and inter-glomerular interactions. The same objective, in the presence of distributed inputs from the motor cortex, implies that the pontine nuclei perform subspace selection. Differences in architecture between the antennal lobe and pontine nuclei are predicted, using our theory, from the labeled line versus distributed organization of their respective inputs. Furthermore, this theory provides an explanation for recent observations of high correlations among granule cells [13]. More generally, our analysis reveals principles that relate the statistical properties of a neural representation to the architectures that optimally transform the representation to facilitate learning.

## Results

The pathways to cerebellum-like structures, such as the mushroom body in the insect olfactory system (Fig. 1a) and the mammalian cerebellum itself (Fig. 1b) are characterized by an initial compression, in which the number of neurons is reduced, followed by an expansion. We model this “bottleneck” motif as a three-layer feedforward neural network (Fig. 1c, see Methods 1). Information flows from *N* input layer neurons to *M* granule cells via a “compression layer” of Nc neurons, each of which samples *L* inputs. Since there are fewer compression layer neurons than inputs the compression ratio *N/N_c_* > 1, while the expansion ratio *M/N_c_* > 1, consistent with the large expansion to granule cells in cerebellum-like structures. In the insect olfactory system (Fig. 1a) tens of olfactory receptor neurons project to an individual glomerulus in the antennal lobe [14]. In the cortico-cerebellar pathway (Fig. 1b) the compression ratio between cortico-pontine projection neurons and neurons in the pontine nuclei is estimated to be between two and ten [8]. In contrast to neurons in expansion layers, which typically emit sparse bursts of action potentials, neurons in compression layers typically have higher firing rates [12,15,16]. For this reason, in our model we consider either linear or rectified linear neurons for the compression layer, while for most of our results we use binary neurons to model the expansion layer.

**Figure 1:**
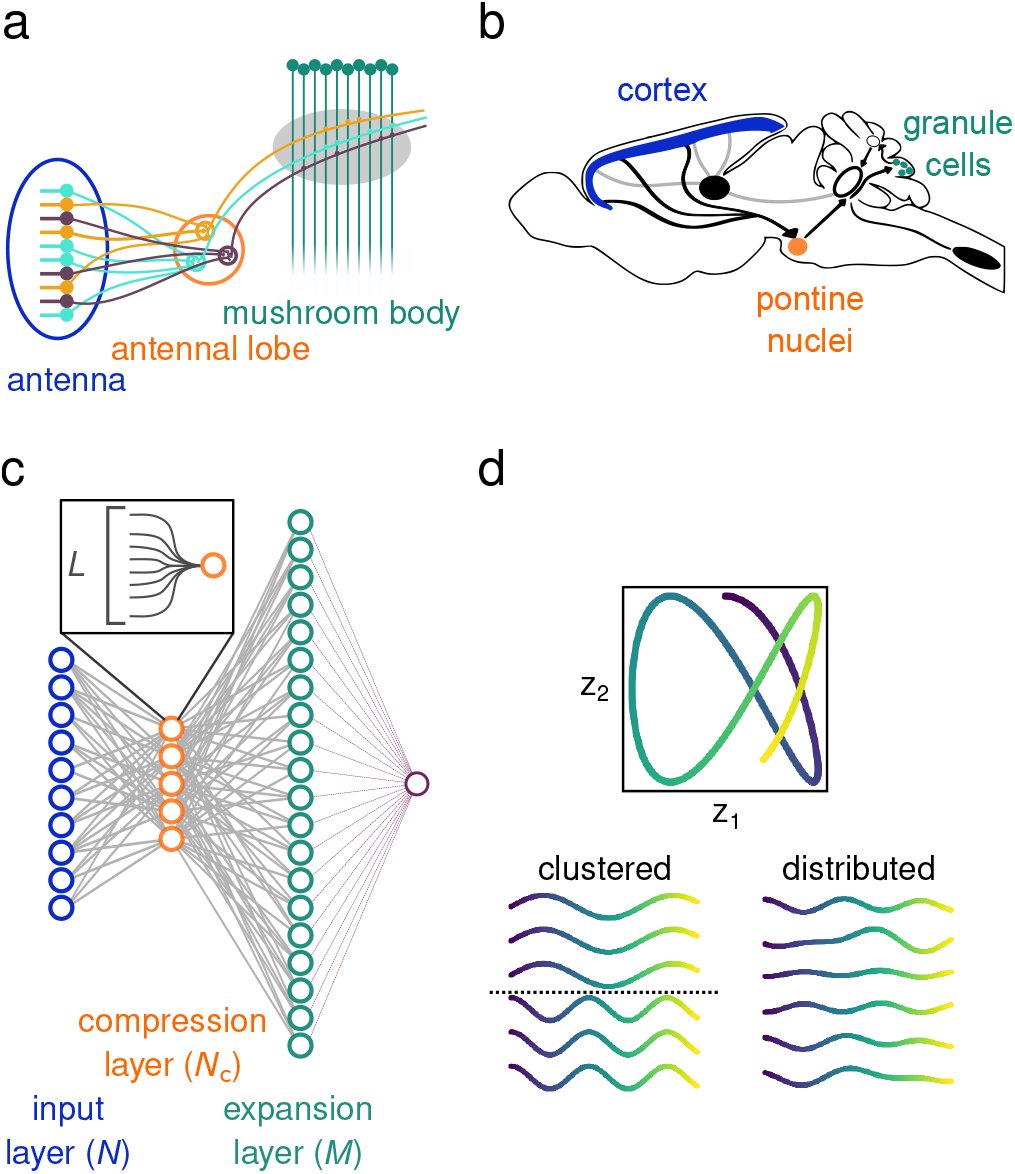
Similar routing architecture to expanded representations. **a:** Schematic of the architecture of the insect olfactory system. **b:** Schematic of the cortico-ponto-cerebellar pathway. **c:** Schematic of the bottleneck model architecture. The input representation is compressed in the compression layer (*N_c_* neurons). Each neuron in the compression layer receives *L* inputs, so that when *L* = *N* the compression is fully-connected. From the compression layer, the representation undergoes a sparse, random and nonlinear expansion to the expansion layer (*M* neurons). Readout weights from the expansion layer (in purple) are adjusted by synaptic plasticity to produce the appropriate output for a specified task. **d:** Example of clustered and distributed input representations. A smooth trajectory in a two-dimensional task space (top) is embedded in an input representation of six neurons (bottom). Bottom left: examples of input neuron responses in a clustered representation (each row is a neuron). The dotted line separates the two clusters. Bottom right: examples of input neuron responses in a distributed representation.

What is the computational role of the compression layer? As a benchmark for comparing different architectures, we begin with the ability of a readout of the expansion layer representation (e.g. a Purkinje cell in the cerebellar cortex) to learn a categorization task in which input layer patterns are associated with positive or negative labels (which could represent positive and negative valences with which conditioned stimuli are associated [6,7]). If the readout is learned using a supervised Hebbian rule, performance on this task increases with the dimension of the expansion layer representation and decreases with its noise strength [7,17] (see Methods 5). We developed a theory describing how these quantities depend on properties of the compression and expansion layer connectivities.

In contrast to previous work [6,7,18], we do not assume that the input patterns are random and uncorrelated. Instead, we define the task as a mapping from patterns in a *D*-dimensional *task subspace* to the labels. The task subspace represents the portion of the input space where inputs relevant to the task tend to lie. For example, in an odor classification task, olfactory sensory neurons of the same type respond similarly, thus defining a subspace in the space of all possible receptor firing rates. In our model, principal components analysis (PCA) performed on the input representation reveals, in the absence of noise, *D* nonzero eigenvalues, with *D* ≪ *N*. These eigenvalues decay (as a power law with exponent *p*; see Methods 3), so that our model exhibits representations similar to those in experiments, for example recordings of motor cortical activity that exhibit such decay [13,19]. We consider two classes of input representations, each of which corresponds to a different organization of selectivities of input neurons to the task variables. In a *clustered* representation, input neurons are organized in distinct groups, each of which is selective to a specific task variable (Fig. 1d, left). Such a representation can arise from a “labeled line” wiring organization and leads to high within-group correlations. In contrast, in a *distributed* representation, each neuron is tuned to different linear combinations of multiple task variables (Fig. 1d, right). As we will show, these two classes of input representation lead to different predictions about the optimal compression architecture, which we relate to differences in anatomy between the antennal lobe and pontine nuclei.

### Selectivity to task-relevant dimensions determines learning performance

We compared the performance of a network without a compression layer, in which input layer neurons are directly and randomly sampled by expansion layer neurons, to two networks with compression: one with random compression weights (representing the synaptic strengths of inputs to the compression layer) and one with learned compression weights. The learned compression weights are trained using error backpropagation [20], under the assumption that the expansion layer connectivity is random and that connections onto the readout neuron are learned using Hebbian plasticity (see Methods 6.5). There is a substantial performance improvement from learning the compression weights, even though the subsequent expansion is fixed and random (Fig. 2a, b). However, the network with random compression performs worse than the network without a compression layer. These results suggest that compression can be highly beneficial, but only if the compression weights are appropriately tuned. If they are not, compression instead degrades performance.

**Figure 2:**
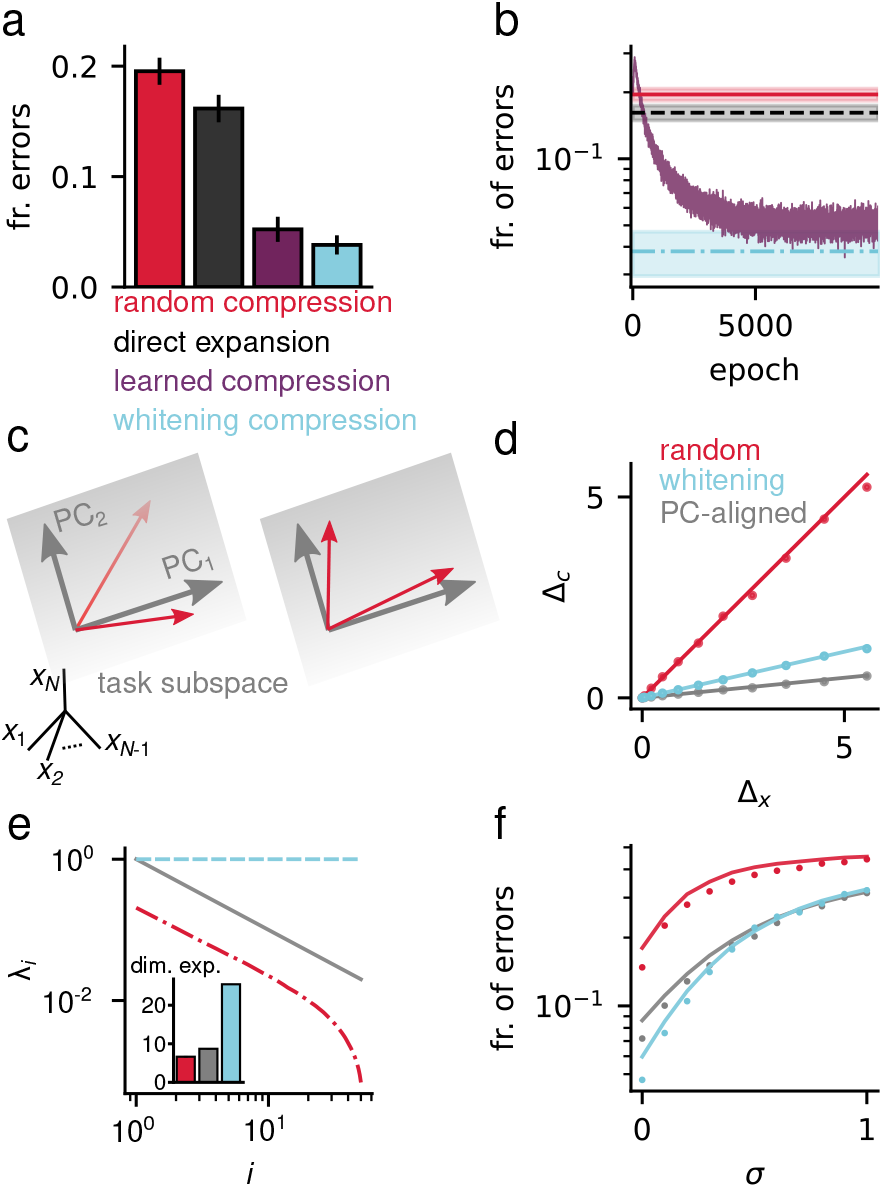
Selectivity to task-relevant dimensions determines learning performance. **a:** Fraction of error of a Hebbian classifier on the random classification task (see Methods 5). *Learned compression* indicates the performance when training via gradient descent has converged (see **b**). The error bars indicate the standard error of the mean across task realizations. **b:** Fraction of errors during training (purple), averaged over task realizations. Color code is the same as in **a**. **c:** Different geometrical arrangements of tuning vectors (red) with respect to the input task-relevant principal components (PCs, gray arrows) and task subspace (gray plane), in the *N*-dimensional input space. Left: tuning vectors do not lie in the task subspace. Right: tuning vectors lie in the task subspace, but are not mutually orthogonal. **d:** Noise strength at the compression layer Δ_*c*_ as a function of the noise at the input layer Δ_*x*_, for random compression (red), whitening compression (light blue), and compression with PC-aligned tuning vectors (gray). Dots are simulation results while solid lines indicate the theoretical prediciton. Noise strength is quantified as the average *cluster size* (see Methods 2). **e:** Spectrum of the task-relevant PCs at the compression layer and corresponding dimension expansion dim(***m*)**/ dim(***x***) (inset), for the same compression strategies as in **d**. **f:** Fraction of errors of a Hebbian classifier on the random classification task, as a function of the input noise standard deviation *σ*. Dots indicates simulations results, lines indicate the theoretical prediction.

To understand the principles underlying these effects, we developed a theory with which we investigate the regimes in which compression is advantageous and explicitly specify the compression weights. In the absence of constraints on compression layer connectivity, and for linear compression layer neurons, our theory demonstrates that expansion layer dimension is maximized and learning performance is increased when compression layer neurons extract the task-relevant principal components of the input representation. As we show later, this simple intuition leads to a number of predictions about the biological implementation of this compression.

For a clustered representation, task-relevant principal components correspond to groups of similarly tuned neurons, while for a distributed representation, they correspond to patterns of activity across the input layer. We represent the preferred stimulus of a compression layer neuron as a “tuning vector” in the *N*-dimensional space of input layer activity. When these vectors lie within the task subspace and are as different as possible across compression layer neurons, the compressed representation is denoised and decorrelated, improving the efficacy of the subsequent random expansion (see Methods 7). Indeed, if a tuning vector lies outside the task subspace, the corresponding compression layer neuron will be in part tuned to task-irrelevant activity, increasing noise (Fig. 2c, d). On the other hand, increased overlap of tuning vectors leads to correlation among compression layer neurons, reducing dimension (Fig. 2c, e).

In addition to alignment with task-relevant principal components, the gain of compression layer neuron tuning affects the dimension of the compressed representation. If the activities of compression layer neurons are scaled so that sub-leading principal components are amplified, dimension is increased. If all principal components are equally strong, the compression layer implements a whitening transformation, which results in the maximum expansion of dimension. Consistent with these results, the performance of a network with whitening compression on a random classification task exceeds that of a network whose compression weights are trained with backpropagation as described above (Fig. 2b). More generally however, we find a trade-off between maximizing dimension and denoising: amplification of sub-leading principal components also causes noise amplification (see Methods 7.2), and whitening ceases to be the best strategy above a certain noise intensity (Fig. 2f). We refer to the network that optimizes the trade-off between dimension and noise as an *optimal compression* network. In the following sections, we describe the architecture of optimal compression networks for two cerebellum-like systems.

So far we assumed that the responses of the compression layer neurons are linear, meaning that the dimension of the compressed representation can be no larger than *D*. However, introducing a nonlinearity at the compression layer can increase the expansion layer dimension, potentially improving input discriminability (Fig. S2; see Methods 5). We therefore ask whether two layers of nonlinear neuronal responses can further improve performance. Surprisingly, in our setting with nonlinear compression followed by random nonlinear expansion, we find they cannot. While nonlinear compression layer neurons do indeed increase dimension, the additional noise introduced by the compression nonlinearity leads to an overall reduced performance. The fact that responses in the antennal lobe and pontine nuclei are substantially denser than those of Kenyon cells or granule cells is consistent with these neurons operating closer to a linear regime, in agreement with our analysis.

### Optimal compression in the insect olfactory system

In the insect olfactory system, olfactory sensory neurons (OSNs) that express the same receptor project to the same olfactory glomerulus in the antennal lobe (Fig. 3a; [14]). Projections from the glomeruli are randomly mixed by Kenyon cells in the mushroom body [22, 23]. We hypothesized that evolutionary and developmental processes optimize the connectivity of the antennal lobe to facilitate a readout of the Kenyon cell representation.

**Figure 3:**
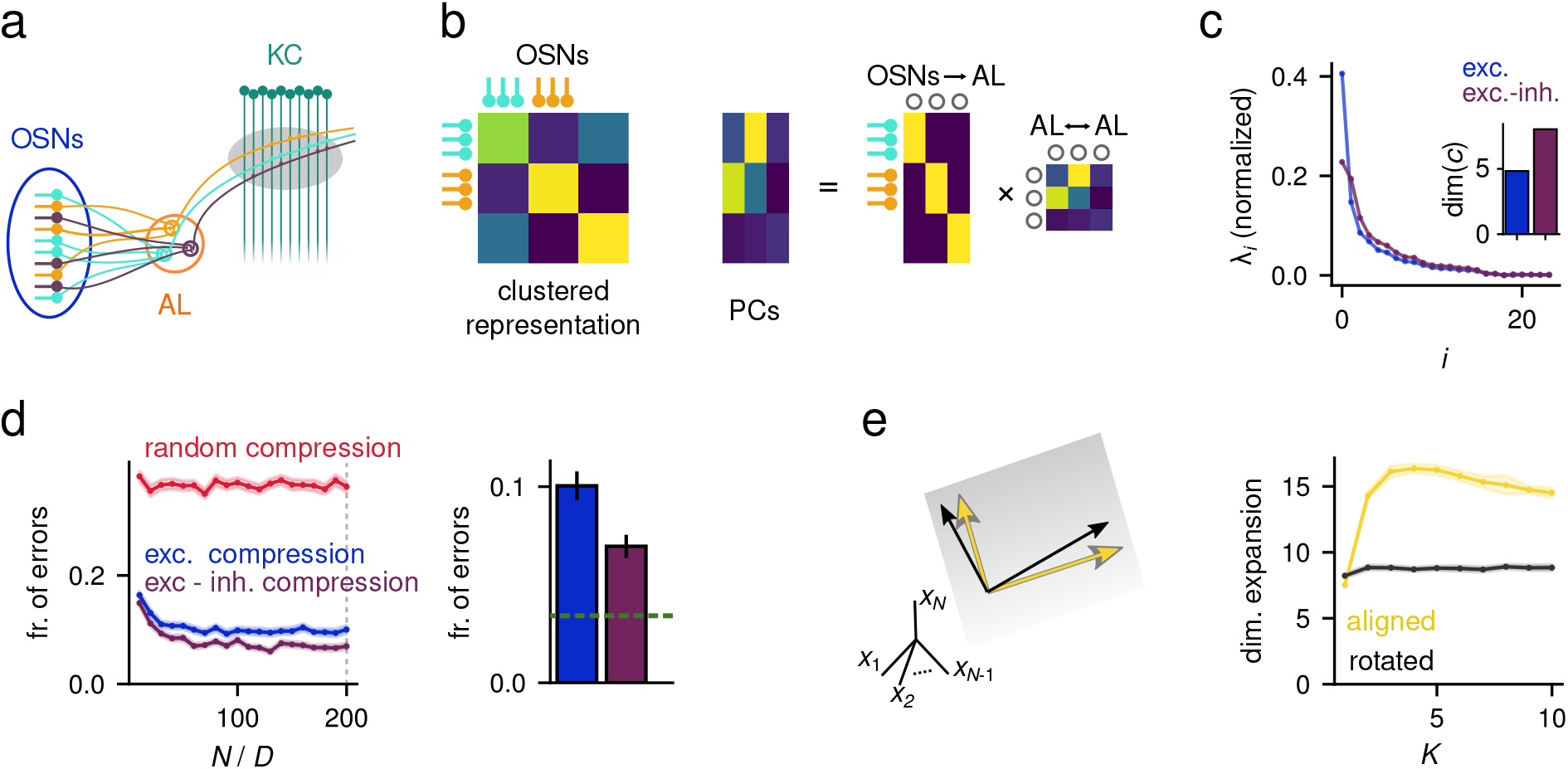
Optimal compression in the insect olfactory system. **a:** Schematic of the insect olfactory system. **b:** Left: Simplified representation of the olfactory receptor neuron covariance matrix. Rows and columns are ordered by receptor type. Center: Matrix whose columns are the principal components of the covariance matrix on the left. This is the transpose of the optimal compression matrix **G**^opt^. Right: Factorization of the principal component matrix in the center panel as a block-diagonal rectangular matrix multiplying a square matrix. The two matrices have a natural biological interpretation as the OSN → AL connectivity and inter-glomeruli interactions, respectively. **c:** Comparison of the compression layer eigenvalue spectrum when inputs are constructed using experimental recordings [21], with and without global inhibition (purple and blue, respectively). The inset shows the corresponding compression layer dimension. **d:** Odor classification performance as a function of the input redundancy *N/D* (left). The bar plot on the right highlights the improvement due to recurrent inhibition (for fixed *N/D* = 200, see vertical dashed line on the left panel) and compares it to the benefit of optimal compression (green dashed line). Shaded region and error bars indicate standard error of the mean. **e:** Left: Geometrical representation of tuning vectors that are aligned (yellow) versus not aligned (black) with principal components, corresponding to clustered and distributed compression layer representations, respectively. Right: Corresponding dimension expansion at the expansion layer plotted against the in-degree of expansion layer neurons *K*.

The OSN representation of odors is clustered, since neurons expressing the same receptor have identical odor tuning. As a result, the covariance matrix of OSN responses is a block matrix, with strong within-block correlations (Fig. 3b, left). According to our theory, in an optimal compression network the compression weights are aligned with the principal components of this covariance matrix. When the responses of OSNs expressing different receptor types are uncorrelated, the compression layer neurons of such a network pool inputs from all OSNs of a particular type, consistent with the anatomical convergence of OSNs to antennal lobe glomeruli. However, OSN responses are correlated, so their principal components have a block structure that is not purely block-diagonal (Fig. 3b, center). We show (see Methods 8) that, in this case, the optimal compression weights can be expressed as a feedforward matrix that represents convergence of OSNs to antennal lobe glomeruli and a square matrix that represents recurrent inter-glomerular interactions (Fig. 3b, right)

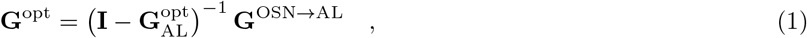

where **G**^opt^ is the optimal compression matrix, 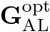 is the matrix of inter-glomeruli effective connectivity that would lead to optimal compression, while **G**^OSN→AL^ contains the OSN to AL compression connectivity.

We reanalyzed experimental recordings of single odor receptors to different odorants [21] to estimate the correlations among OSN types and found that they are more positive than expected by chance (Fig. S3). We show analytically that when these correlations are uniformly positive, global lateral inhibition across antennal lobe glomeruli is sufficient for optimal compression (see Methods 8). Consistent with this result and with studies that propose inter-glomerular interactions perform pattern decorrelation and normalization [10,11,24], global inhibition considerably increases the dimension of the antennal lobe representation when using the recorded responses as input to our model (Fig. 3c). This increase in dimension leads to improved performance in an odor classification task (Fig. 3d). However, correlations are not uniformly positive, so a network with global inhibition does not reach the performance of a network in which the antennal lobe representation is perfectly decorrelated (Fig. 3d, right), since specific lateral interactions among glomeruli further increase dimension in this case. Future studies should analyze whether the specific structure of lateral connectivity in the antennal lobe is consistent with this role (see Discussion).

In contrast to networks with specific convergence of OSN types onto glomeruli, a network in which OSNs are randomly mixed in the antennal lobe performs poorly (Fig. 3d, left). It may seem counterintuitive that such convergence is needed for optimal performance when antennal lobe responses are subsequently randomly mixed by Kenyon cells. Our theory illustrates that this difference is a consequence of both denoising and maximization of dimension. When OSN responses are noisy, pooling OSNs of the same type reduces noise by a factor *N/D* compared to random compression (see Methods 7.2). Even in the absence of noise, the dimension of the compression layer is higher for clustered compression than for random compression. This arises from the random distortion of the input layer representation introduced by random compression weights. This distortion can only be avoided by ensuring weights onto compression layer neurons are orthogonal, a more stringent requirement that cannot be assured by independent random sampling of inputs (see Methods 7.1). In fact, if input to compression layer weights are constrained to be excitatory, the observed OSN convergence is the only possible weight matrix that has this property.

While the above benefits of convergence can be understood in terms of a linear mapping from input to compression layer, an additional and more subtle benefit of clustered compressed representations arises from the sparseness of inputs to the nonlinear expansion layer. Even when both representations are constructed with orthogonal compression weights, a sparse expansion leads to a higher dimension for a clustered representation than for a distributed one (Fig. 3e). This result arises from the ability of sparsely connected expansion layer neurons to select different principal components. If these neurons sample from a distributed representation, their input is always dominated by leading components and sub-leading ones are discarded, decreasing dimension. In contrast, if they sample only a few inputs from a clustered representation, some neurons mix sub-leading components, increasing dimension. In total, our theory reveals analytically that OSN convergence and inter-glomerular inhibition can be viewed as consequences of optimizing the Kenyon cell representation to facilitate downstream learning.

### Optimal compression in the cortico-cerebellar pathway

In the cortico-cerebellar pathway, inputs from motor cortex are relayed to cerebellar granule cells via a compressed representation in the pontine nuclei. The motor cortex representation is low-dimensional and distributed (Fig. 4a; [25]), as opposed to the clustered representation of inputs to the antennal lobe. Its principal components contain both positive and negative elements, but since direct cortico-pontine projection are excitatory, effective input layer to compression layer weights can align with these components only if effectively inhibitory weights result from either di-synaptic feedforward inhibition or recurrent inhibition. Unlike the antennal lobe, the pontine nuclei lack strong lateral inhibition. In rodents, lateral inhibition seems to be completely absent, while for primates and larger mammals, it appears to play only a limited role [8].

**Figure 4:**
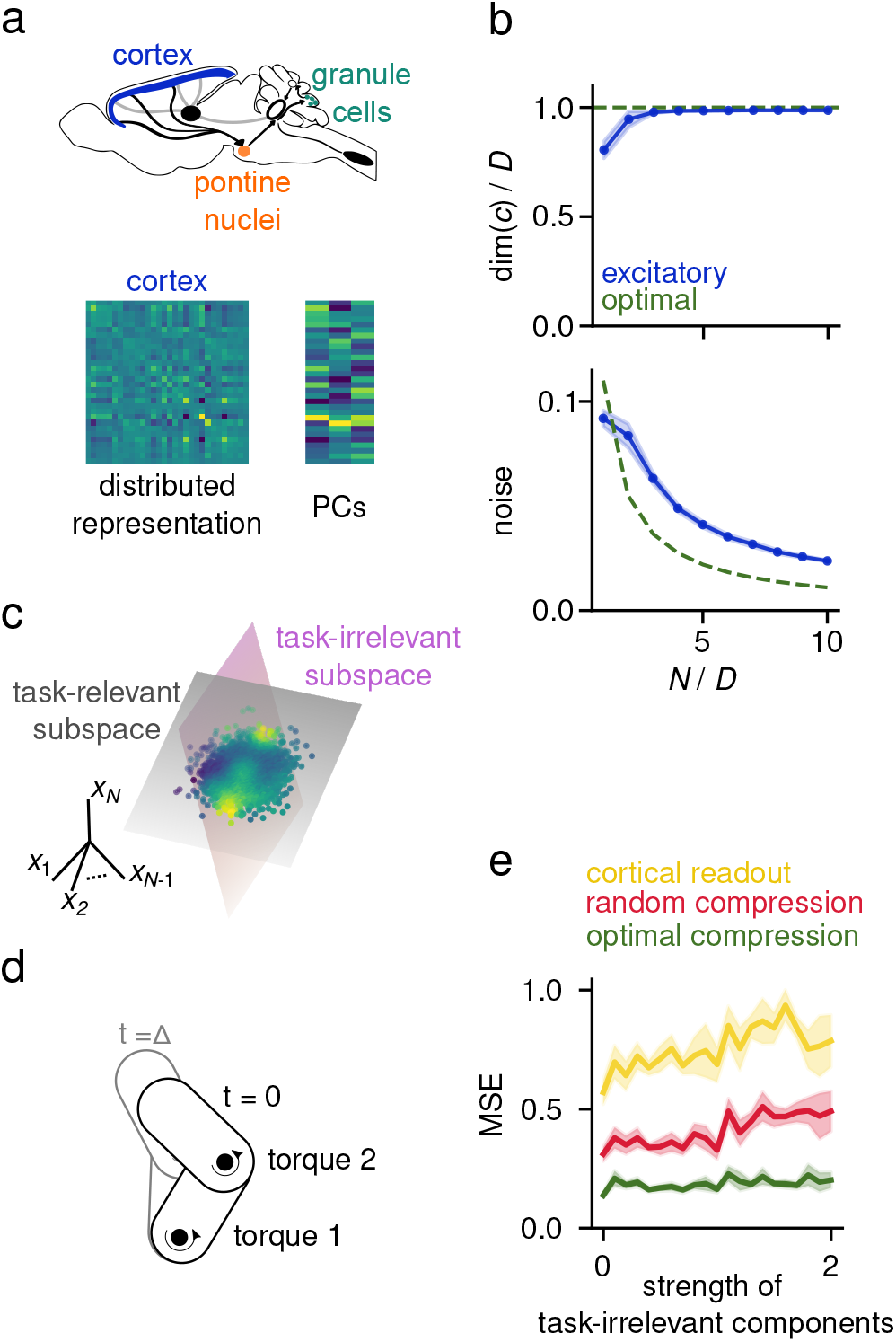
Optimal compression in the cortico-cerebellar pathway. **a:** Top: Schematic of the cortico-ponto-cerebellar pathway. Bottom left: Example of covariance matrix for a distributed representation. Bottom right: Matrix of corresponding principal components (one column per component). Since principal components have positive and negative entries, optimal compression weights requires both excitatory and inhibitory connections. **b:** Sign-constrained compression. Results of gradient descent training when the compression weights were constrained to be positive (blue), compared to optimal compression (green), for a distributed input representation. The training objective was to maximize dimension at the compression layer while simultaneously minimizing noise (see Methods 6.5). Top: dimension of the compression layer representation, as a function of the input redundancy *N/D.* Bottom: Same as top, but for noise strength Δ_*c*_ in the compression layer. Shading indicates standard deviation across input realizations and weight initializations. Parameters: *D* = 10, input noise strength *σ_x_* = 0.2, *p* =1, *N_c_* = 10. **c:** Illustration of a continuously varying target in the cortical representation. The high-dimensional cortical representation consists of orthogonal task-relevant and task-irrelevant subspaces. The cerebellum learns to map cortical activations (dots) to an output (color code) via a smooth nonlinear function. **d:** Schematic of two-joint arm task. Given the state of the arm, defined by two joint angles and two angular velocities, and the torque applied to both joints, the system predicts the state of the arm at a time point Δ time units in the future. **e:** Mean-squared error (MSE) of the model in predicting the future state of the arm, plotted against the strength of task-irrelevant activity. “Cortical readout” indicates the performance when reading out directly from the input representation, which can be considered a performance baseline. Network parameters: *M* = 1000, *N* = 500, *N_c_* = 100, *L* = 50. Task parameters: Δ = 0.1*s*

We therefore asked whether purely excitatory compression can approximate optimal compression for a distributed representation. We use gradient descent to train excitatory feedforward compression weights, with the objective of maximizing dimension and minimizing noise at the compression layer (see Methods 6.5). Surprisingly, when the input representation is distributed and redundant (*N* ≫ *D*), purely excitatory compression leads to a compression layer dimension comparable to the one obtained via optimal compression (Fig. 4b, top). This is because in this scenario it is possible, with high probability, to find input neurons that encode each dimension of the task subspace. Projections from these neurons to the compression layer are sufficient to avoid a decrease in dimension. While purely excitatory compression is less effective in filtering out input noise, this limitation can be compensated by increasing the input redundancy (i.e. making *N/D* larger, see Fig. 4b, bottom). Thus, even in the absence of lateral inhibition, excitatory cortico-pontine projections can be adjusted to maximize classification performance at the Purkinje cell readout (Fig. S4a, b). This result stands in contrast to our previous conclusion for systems with clustered input representations, such as the antennal lobe, for which lateral inhibition is necessary to decorrelate the representation, and for which increasing the input redundancy is not a viable solution (Fig. 3dc, d and Fig. S4c). This difference in architecture can therefore be explained in terms of the distributed nature of the motor representation.

So far, our theory and results have focused on optimizing performance for a classification task. Some of the tasks that the cerebellum is involved in, such as eye-blink conditioning or timing tasks, may be reasonably interpreted in this way, but others may not be. An influential hypothesis of cerebellar function is that the cerebellum predicts the sensory consequences of motor commands, implementing a so-called forward model [26]. In this view, the cerebellum integrates representations of the current motor command and sensory state to estimate future sensory states (Fig. S8). We cast the problem of learning a forward model as a nonlinear regression task, assigning each point in the task subspace (representing the combination of motor command ***u*** and sensory state **s**) a predicted sensory state *s’* = ***s*** + *f* (***s***, ***u***) (Fig. 4c, see also Methods 10). In this case, the goal of cerebellar Purkinje cells is to learn the nonlinear function *f* (***s***, ***u***). We consider a planar two link arm model, characterized by two joints at which torques can be applied (Fig. 4d). Optimal compression leads to substantially better performance than random compression when learning a forward model for this system, showing that the benefits we have described are not specific to discrete classification tasks (Fig. 4e).

### Bottleneck architecture is more efficient than a single layer network

Could we attain the benefits of optimal compression without requiring two layers of processing from input to expanded representation? Since optimal compression, as we have defined it, involves a linear transformation, it is indeed possible to generate an equivalent single-layer expansion by setting the expansion weights equal to the product of the optimal compression and random expansion weights. What then is the advantage of performing these operations in two distinct steps? The answer is a consequence of the sparsity of the expansion layer weights.

For the expansion layer to implement both optimal compression and dimensional expansion, its neurons must be equipped with a local decorrelation mechanism across their afferent synapses (Fig. 5a, inset). However, due to their sparse connectivity, each expansion layer neuron receives input only from a subset of input neurons. Minimizing correlations of this sub-sampled representation is unlikely to lead to decorrelation of the full representation. As a result, the benefit of adding local whitening to the single-layer expansion architecture is limited (Fig. 5b) if the in-degree of the expansion layer is small and neurons are not permitted to use non-local information (information about neurons to which they are not connected) in order to set synaptic weights.

**Figure 5:**
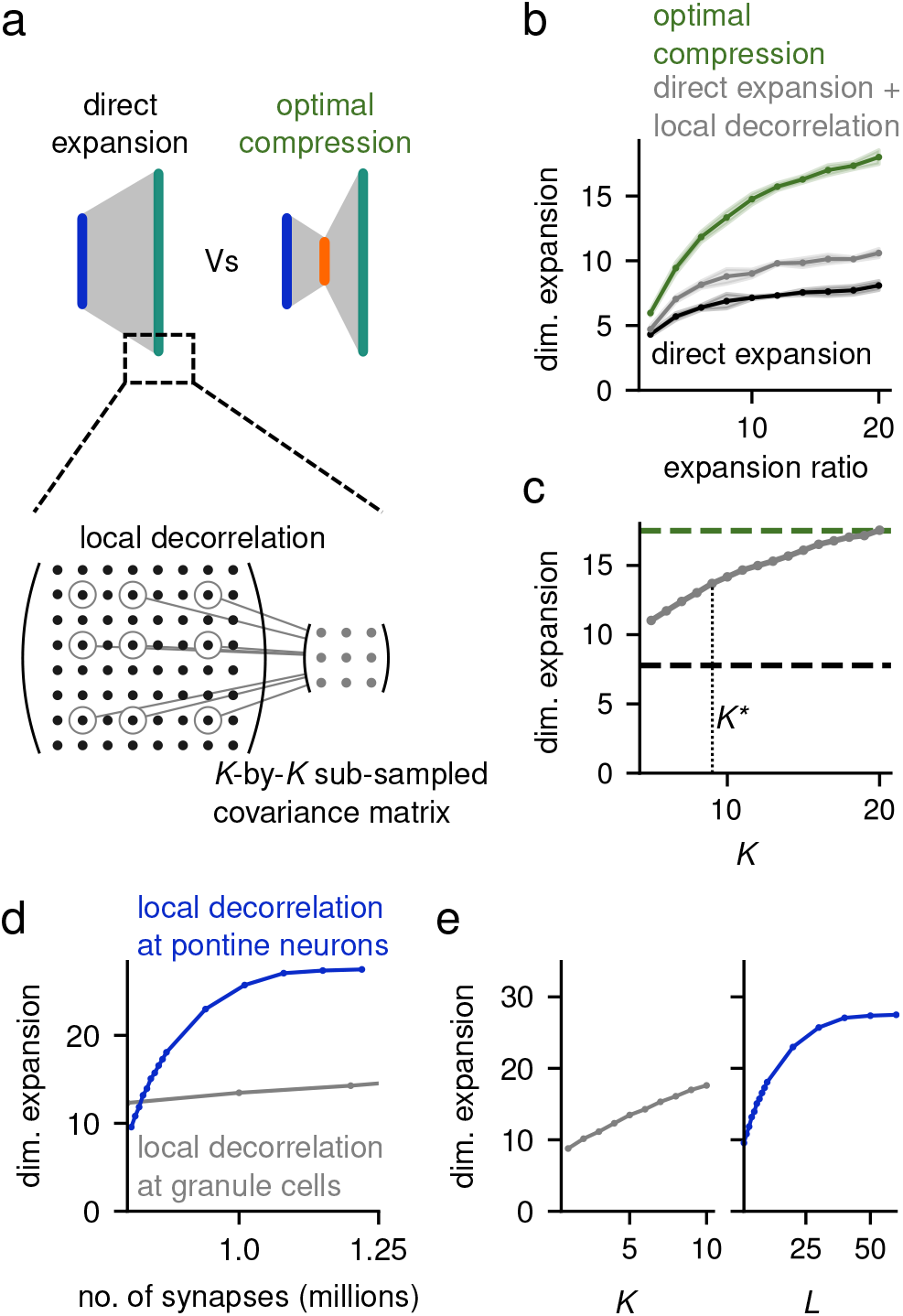
Bottleneck architecture is more efficient than a single layer network. **a:** Direct expansion (left) and optimal compression (right) architectures. Inset: Illustration of local decorrelation at granule cells in the direct expansion architecture. With sparse connectivity, each expansion layer neuron only has access to a sub-sampled version of the full input covariance matrix (3-by-3 in the illustration). We implemented local decorrelation by having each expansion layer neuron decorrelating this sub-sampled representation and nonlinearly mix the resulting representation. **b:** Dimension expansion dim(***m***)/dim(***x***) for different network architectures plotted against the expansion ratio. Local decorrelation yields only a small improvement. **c:** Dimension expansion yielded by local decorrelation for increasing *K*. The total number of synapses quickly increases with *K*, and *K*\* (dotted line) indicates the value at which it becomes larger than the number of synapses needed with optimal compression. **d:** Dimension expansion plotted against the total number of synapses used in the bottleneck architecture with local decorrelation at the compression layer (blue) and for a direct expansion architecture with local decorrelation at the expansion layer (gray). **e:** Same as **d**, but plotted against the granule cell indegree *K* for the direct expansion architecture (left) and against the pontine in-degree *L* for the bottleneck architecture (right).

If we increase the expansion layer in-degree in the model, local decorrelation better approximates optimal compression (Fig. 5c). However, the total number of synapses necessary to implement this architecture is larger than that required for optimal compression. The wiring cost of performing local decorrelation at the expansion layer is particularly high when considering parameters consistent with cerebellar cortex, i.e. *M* ≃ 200, 000 and *N_c_* ≃ 7000. In this situation, it is much more efficient to perform local decorrelation at the cortico-pontine connections than at the granule cell layer (Fig. 5d,e). With local decorrelation at the cortico-pontine connections, dimension expansion saturates when the pontine in-degree *L* is around 40 (Fig. 5e. right), totaling slightly more than a million synapses. In contrast, we expect that to achieve the same expansion of dimension via local decorrelation at the granule cell layer, *K* would need to be between 20 and 40 (extrapolating from Fig. 5e, left), for which the total number of synapses would be between 4 and 8 million. We reach a similar conclusion when considering parameters consistent with the insect olfactory system (Fig. S5). In summary, our results show that a dedicated compression layer provides an efficient implementation of this computation, both in terms of number of synapses and wiring complexity.

### Optimal in-degree of learned cortico-pontine compression

Activity in motor cortex is task-dependent and exhibits steady drift in the neurons representing a stable latent dynamics [27]. Unlike the genetically determined, clustered representation of olfactory sensory neurons, such activity thus has a covariance structure that changes over time. We therefore extended our theory from the case of fixed covariance to one that must be learned through experience-dependent synaptic plasticity.

Hebbian plasticity rules are a natural candidate for learning compression weights, because they enable downstream neurons to extract the leading principal component of upstream population activity [28–30]. In many models, recurrent inhibitory interactions among downstream neurons are introduced to ensure that each neuron extracts a different principal component. Due to the lack of inhibition in the pontine nuclei, we asked whether sparsity of compression connectivity instead can introduce the necessary diversity among pontine neuron afferents to achieve high task performance.

When compression weights are set using Hebbian plasticity (see Methods 6.6), the performance of a Hebbian classifier trained on the expansion layer representation has a non-monotonic dependence on *L*, the in-degree of the pontine neurons. The performance is poor for very small *L*, increases quickly, and finally decays slowly as *L* becomes very large (Fig. 6a, left). This behavior is a result of the trade-off between denoising and dimension. Noise strength at the expansion layer decays with *L*, because increased *L* permits more accurate estimation of leading principal components (Fig. 6b, top right). On the other hand, dimension decreases with *L*, since as *L* increases, compression layer neurons estimate similar components (Fig. 6b, bottom right). The value *L*\* that yields the best performance lies between ten and a hundred incoming inputs. *L*\* is only weakly affected by architectural parameters such as the number of input neurons *N* or expansion layer neurons *M* (Fig. S6c). Instead, it depends on features of the input representation, such as its dimension and the noise strength, with stronger noise and lower-dimensional representations favoring large in-degrees (Fig. 6b,c).

**Figure 6:**
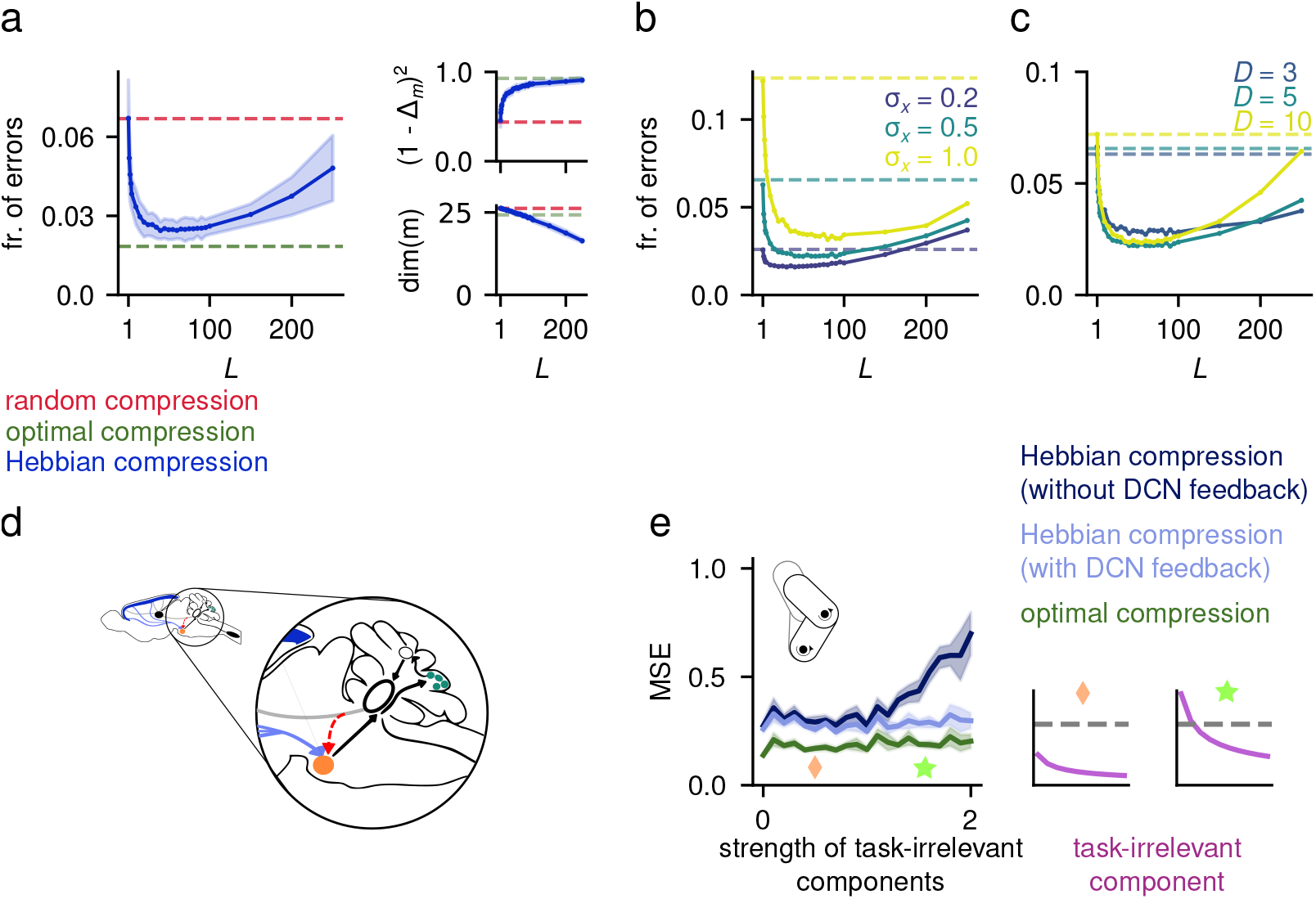
Biologically plausible learned compression. **a:** Fraction of errors (left) of a Hebbian classifier reading out from the expansion layer representation for random, PCA and Hebbian compression, as a function of the compression layer in-degree *L*. On the right, we show the dimension (bottom) and noise (top) contributions to the classifier SNR, which result in the non-monotonic behavior of the performance. **b:** Effect of noise strength on the network performance. **c:** Effect of input dimension on the network performance. For both panels **b** and **c**, the solid line indicate Hebbian compression, while the dashed lines indicate random compression. In panels **a**, **b** and **c** we used *N* = 500, *N_c_* = 250, *M* = 1000, *D* = 5, *p* = 0.1 and *σ_x_* = 0.5 unless otherwise stated. **d:** Illustration of the DCN-pontine feedback (in red). **e:** Feedback from DCN improves selection of the task-relevant dimensions. Mean-squared error (MSE) of the bottleneck network on the two-joint arm forward model, as in Fig. 4e. Inset: variance explained by task-irrelevant components (violet), in decreasing order, for two example values of the task-irrelevant component strength (green star and orange diamond). For comparison, gray dashed line indicates the variance explained by the leading task-relevant component. Network parameters: *M* = 1000, *N* = 500, *N_c_* = 100, *L* = 50. Task parameters: Δ = 0.1*s*.

Thus, in the absence of recurrent inhibition in the compression layer, an intermediate in-degree leads to the best performance. This is not only true for random classification tasks, but also for nonlinear regression (Fig. S7b). Given the typically low-dimensional motor cortex representation, we predict that the optimal in-degree of rodent pontine nuclei neurons should be between ten and a hundred. To our knowledge, this in-degree has not been measured, but the large dendritic arbor of these neurons [31], suggests that it is at least much larger than the in-degree of granule cells.

We also tested whether further improvement could be achieved when we include recurrent inhibition using a recent model that implements a combination of Hebbian and anti-Hebbian plasticity rules [30, 32]. After learning, the average principal component overlaps of sub-leading components decay more slowly than without recurrent inhibition (Fig. S6e), improving performance (Fig. S6f). As a consequence, we predict that species with more recurrent inhibition in the pontine nuclei may exhibit larger pontine in-degree.

### Feedback from the deep cerebellar nuclei improves selection of task-relevant dimensions

One limitation of tuning the compression weights using Hebbian plasticity is that by construction it is an unsupervised method, meaning that it extracts leading principal components, but not necessarily only task-relevant ones. This might not present a problem when the input is only corrupted with independent, high-dimensional random noise, since in this case the PCA spectrum is likely to be dominated by signal components. However, it will reduce the performance when leading components are task-irrelevant. The anatomy of the cortico-cerebellar system suggests a solution to this problem: in addition to cortical input, the pontine nuclei also receive feedback from the deep cerebellar nuclei (DCN), the output structure of the cerebellum [31] (Fig. 6d). Previous theories have largely ignored these connections. We provide a novel interpretation of this motif and suggest that it provides a supervisory signal that aids identification of task-relevant inputs.

To test this hypothesis, we extended our model to include feedback of the target output to pontine neurons. This feedback is used solely as a supervisory signal for synaptic plasticity, i.e. it is added as a regular input to the learning rule, but it does not affect the network dynamics (see Methods 9.1). Specifically, we considered a variant of Oja’s rule [28] that is augmented with supervisory feedback from the DCN, and show that such plasticity is biased towards components of the input which correlate with the target and are therefore likely task-relevant (see Methods 9.1).

We tested this mechanism in the two-joint arm forward model task considered above. When the task-irrelevant components of the input are weak, Hebbian compression performs well, both with and without feedback (Fig. 6e). In contrast, when task-irrelevant components are stronger than task-relevant ones (Fig. 6e, inset), the performance of Hebbian compression in the absence of feedback quickly degrades, as compression layer neurons learn to extract task-irrelevant components. However, in the presence of supervisory DCN feedback, compression layer neurons always extract only task-relevant input components, resulting in a stable performance (Fig. 6e). In summary, our results support a novel functional role for DCN input to pontine neurons: enabling the extraction of task-relevant, but sub-leading, input components.

### Hebbian compression explains correlation and selectivity of granule cells

Recent recordings have shown that cerebellar granule cells in mice exhibit high selectivity to task variables and strong correlations in activity, both with each other and with cortical neurons [13]. Both findings are, at first sight, at odds with theories of random mixing [6, 7, 33] which predict that granule cells are decorrelated and encode nonlinear combinations of task variables. We show that optimal compression in the cortico-pontine pathway provides an alternative explanation for these experimental findings that preserves mixing in the granule cell layer.

We developed a model based on simultaneous two-photon calcium recordings of layer-5 pyramidal cells in motor cortex and cerebellar granule cells (Fig. 7a; [13]). During recording sessions, mice performed a skilled forelimb task (see Methods 11) that required them to move a joystick in a L-shaped trajectory, turning either to the left or to the right. We used recorded calcium traces of layer-5 pyramidal cells as inputs ***x*** to the cortico-ponto-cerebellar model described above. In the model, cortico-pontine synapses undergo unsupervised learning via Hebbian plasticity. Similar to [13], we modeled unrecorded neurons by including an unobserved layer-5 population and associated pontine subpopulation. The latter projects to both the observed granule cell layer and other unobserved granule cells that we do not include in the model (Fig. 7b).

**Figure 7:**
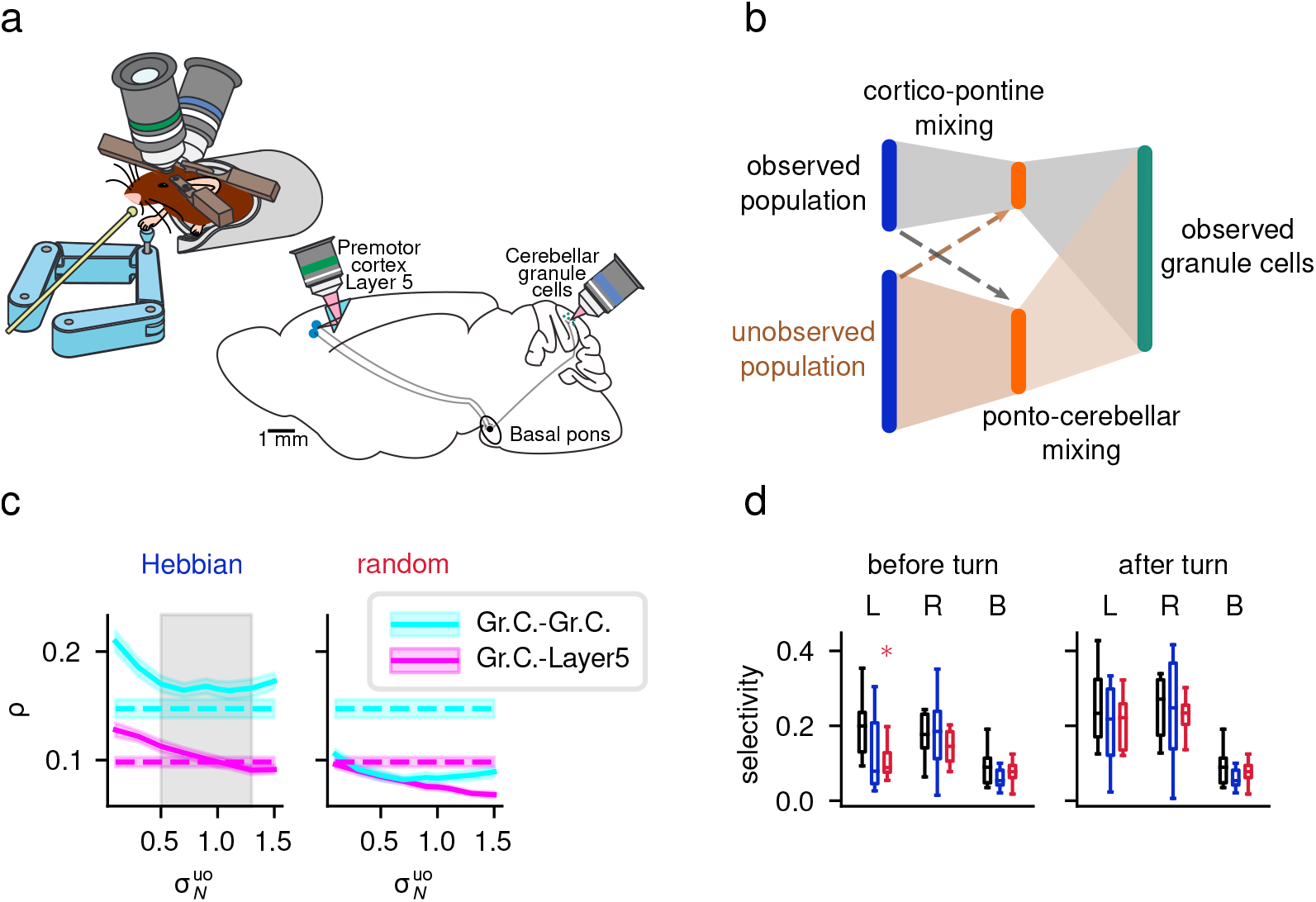
Bottleneck model explains correlations and selectivity of recorded granule cells. **a:** Illustration of the experimental design of [13]. Mice performed a forelimb control task (left) while layer-5 pyramidal neurons and cerebellar granule cells were recorded simultaneously using two-photon calcium imaging (right). **b:** Schematic illustrating how the bottleneck model is extended to reproduce the data. The dashed line indicates little or no mixing in the cortico-pontine pathway, while the shaded areas indicate strong mixing in the pontocerebellar pathway. **c:** Layer 5-granule cell (magenta) and granule cell-granule cell (cyan) correlations, for Hebbian (left) and random (right) compression strategies. Mean correlations across neurons are averaged across animals and plotted against 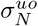, the noise strength in the unobserved population. Colored shaded area indicates standard error of the mean, computed across animals. Gray shaded area indicates the region in which correlations in the model are not statistically different from those in the data for both areas (*p* > 0.05, Wilcoxon signed-rank test). For this panel, the signal strength of the unobserved population is 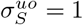. **d:** Average selectivity to left (L) and right (R) turns of the joystick, or to turns without direction preference (B), for granule cells in the data (black) and models (Hebbian: blue; random: red). Selectivity is measured separately for the time-window before the turn (left) and after the turn (right). Boxes indicate 25th and 75th percentiles across mice, while whiskers indicate the full range of the mean selectivities across mice. Asterisks indicate cases in which model and data are not compatible, color coded according to the compression type. For panel **d**, 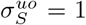 and 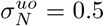 for the Hebbian model and 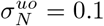 for the random model. For all panels, *N_c_* = *N*/2, *M* = 10*N*, *N_uo_* = 2*N*, *f* = 0.1, *L* = 20.

Since we do not have access to the unobserved population, we introduce two model parameters 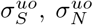, which control the strength of task-relevant (signal) and task-irrelevant (noise) components of the unobserved cortical population (see Methods 11). We systematically varied both parameters and measured average correlations in the model, both among granule cells and between granule and layer-5 cells. The model and data are compatible with respect to both measures, provided that task-irrelevant activity in the unobserved cortical population is strong enough (Fig. 7c, left). Notably, a model with random, non-plastic compression weights is not compatible with the data and exhibits lower correlations even for very small 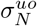 (Fig. 7c, right).

We also quantified the selectivity of granule cell subpopulations responsive to left and right turns of the joystick, or responsive to both directions, before and after the turn ([13], see Methods 11). We fix the parameters of the unobserved population to those yielding the best match of the granule cell correlations, with no further tuning. The model accounts for the selectivities observed in the data (Fig. 7d). A model without Hebbian compression can also explain the selectivity profile, but only if task-irrelevant activity in the unobserved population is assumed to be extremely weak.

Our analysis shows that the results of [13] are consistent with granule cells responding to mixtures of mossy fiber activity. Due to Hebbian plasticity, neurons in the pontine nuclei filter out task-irrelevant activity, becoming more selective to task variables than what would be expected from random compression and forming a lower-dimensional task representation. This decrease in dimension is not detrimental, but rather a consequence of discarding high-dimensional, task-irrelevant activity and preserving task-relevant activity. Since the latter is lowdimensional, random mixing at the granule cell layer yields only a moderate dimensional expansion and high correlations. Altogether, this analysis shows that for low-dimensional tasks, unsupervised learning at the corticopontine synapses leads to a selective and highly correlated granule cell representation, despite mixing of distinct mossy fiber inputs. This result supports structured, as opposed to random, compression at the pontine nuclei.

## Discussion

Our results demonstrate that specialized processing in “bottleneck” structures presynaptic to granule-like expansion layers substantially improves the quality of expanded representations. This two-stage architecture, with a structured bottleneck followed by a disordered expansion, is also more efficient, in terms of total number of synapses, than a single-stage architecture. In the insect olfactory system, our theory shows that the glomerular organization of the antennal lobe is optimized to maximize the dimensionality of the Kenyon cell representation for clustered input. In the cortico-cerebellar pathway, it shows that Hebbian learning of excitatory cortical inputs improves performance for distributed input and provides a novel prediction for the in-degree of pontine neurons as well as the role of feedback from the deep cerebellar nuclei. It also accounts for the magnitude of correlations among simultaneously recorded granule cells and motor cortical neurons, arguing that for tasks with low-dimensional input representations, low-dimensional granule cell representations are optimal.

### Other pathways to the cerebellum and other cerebellum-like structures

We focused on the cortico-ponto-cerebellar pathway and the insect olfactory system due to the ability to characterize the statistics of their inputs, but other cerebellar regions and cerebellum-like structures also exhibit bottleneck architectures. In addition to the cerebrocerebellum that is targeted by pontine inputs, the spinocerebellum is innervated by inputs from the spinal cord that integrate proprioceptive information from skeletal muscles and joints [34]. Characterizing the statistics of this ensemble of proprioceptive inputs to predict the optimal organization and connectivity of this pathway is an interesting direction for future research.

In electric fish, the nucleus praeeminentialis (PE) is a major source of input to the electrosensory lobe [2]. PE receives input from the midbrain and cerebellum, while sending output solely to the electrosensory lobe. Interestingly, PE also receives inputs from the electrosensory lateral line lobe (ELL) neurons, a pathway analogous to DCN-pontine nuclei feedback connections. This suggests that our hypothesized supervisory role of DCN-pontine nuclei feedback could be an instance of a more general motif across cerebellum-like structures.

### Response properties of compression layer neurons

While in previous work [13] pontine neurons have been modeled as binary, here we consider linear neurons, which we argue is more consistent with the graded firing rates they exhibit. Indeed, pontine neurons have higher firing rates and denser responses than cerebellar granule cells [12,15,16]. The latter are modeled here and in previous work as binary units, motivated by their tendency to reliably respond with sparse spikes or bursts to combinations of inputs [15]. Similar arguments are valid for the insect olfactory system, when comparing projection neurons, which have denser responses, to Kenyon cells [16].

We tested that our results are robust when pontine neurons are modeled using rectified linear units (ReLUs) and showed that the introduction of nonlinear responses at the level of the compression layer does not improve the performance of the bottleneck network. This reflects that a linear transformation is well-suited to maximize the performance of the subsequent random expansion. However, it is also possible that, for specific input statistics, nonlinear compression layer neurons lead to an improvement. Furthermore, non-random expansion architectures, such as deep networks, can benefit substantially from multiple nonlinear layers [1].

### Feedforward and lateral inhibition

In most mammalian brain areas, long-range projections are predominantly excitatory. This is true of corticopontine projections from layer-5 pyramidal cells and the cholinergic projections of OSNs to the antennal lobe [35]. When the input representation is clustered into groups of neurons that exhibit high correlations within each cluster and are uncorrelated across clusters, we showed that convergence of projections from each cluster, such as the glomerular organization of the antennal lobe, is optimal. When correlations between clusters exist, we showed that either di-synaptic feedforward or lateral inhibition is necessary to maximize performance.

In the antennal lobe, both types of inhibition are present. However, di-synaptic inhibition is believed to largely mediate interactions among different glomeruli [10], suggesting that lateral inhibition dominates. We showed that global lateral inhibition is sufficient to effectively denoise and decorrelate OSNs whose response properties are constrained by experimental data [21]. It is an interesting future direction to investigate whether the pattern of correlations across specific pairs of glomeruli are reflected in lateral antennal lobe connections [36]. However, such a prediction requires accurate estimation of this correlation pattern over the distribution of natural odor statistics, which may not be reflected in existing datasets [21].

While inhibitory di-synaptic pathways to the pontine nuclei do exist [8,12], our results suggest that purely excitatory compression weights can perform near-optimally when the input representation is redundant and distributed, rather than clustered. However, we find that lateral inhibition might play a role in learning. Lateral inhibition promotes competition to ensure heterogeneous responses even when compression layer neurons share many inputs [30]. While lateral inhibition is almost absent from the pontine nuclei in rodents, its presence increases in larger mammals, such as cats and primates [8]. This suggests that in species where lateral inhibition is more abundant, pontine neurons may be more specifically tuned to task-relevant input dimensions and may exhibit larger in-degrees.

### Cortico-pontine learning and topographical organization of the pontine nuclei

Our theory highlights the importance of plasticity at cortico-pontine synapses, thanks to which pontine neurons select task-relevant subspaces within the cortical input space. This could support, for example, distinguishing between movement preparation and execution [37] or compensating for representational drift in motor cortex [27]. We also showed that such subspace selection can be further improved by supervisory feedback from the DCN. Each pontine neuron approximately extracts task-relevant principal components, which requires a large number of pontine neurons to achieve good performance (Fig. S6a,b). This result provides a motivation for the smaller compression ratio observed in the pontine nuclei compared to the antennal lobe: when subspace selection is imperfect, it must by compensated by a larger number of neurons in the compression layer. Another possibility is that part of the learning process that enables subspace selection is carried out by layer-5 pyramidal cells. These neurons receive a large number of inputs from other cortical neurons and therefore are capable of performing some of the computations we have described.

At a larger scale, the pontine nuclei exhibit a topographical organization, perhaps genetically determined, that largely reflects the cortical organization [38,39]. For example, motor cortical neurons responsible for different body parts project to distinct regions of the pontine nuclei. Moreover, there is evidence of convergence of motor and somatosensory cortical neurons coding for the same body part onto neighboring pontine regions [40]. It is therefore likely that both hard-coded connectivity and experience-dependent Hebbian plasticity control the compression statistics.

### Random mixing and correlations in low-dimensional tasks

Our theory is consistent with data collected using simultaneous two-photon imaging from layer-5 pyramidal cells in motor cortex and cerebellar granule cells [13]. We showed that the level of correlations and selectivity of granule cells can be explained if cortico-pontine connections are tuned to task-relevant dimensions, but not if they are fixed and random. A previous theory proposed that the data could be accounted for by a model in which, during the course of learning, mixing in the granule cell layer is reduced and for each granule cell a single mossy fiber input comes to dominate its response [13]. Our model preserves mixing in granule cells, a feature thought to be crucial for the computations performed by the cerebellar cortex [3], and instead emphasizes the role of low-dimensional granule cell representations when animals are engaged in behaviors with low-dimensional structure. Such an interpretation may generally account for recordings of granule cells that exhibit low dimensionality and suggests the importance of complex behavioral tasks or multiple behaviors to probe the computations supported by these neurons [41].

## Acknowledgements

We would like to thank Marjorie Xie, Adam Hantman, Britton Sauerbrei, Jonathan Kadmon, and Rick Warren for helpful discussions and comments. We would also like to thank Larry Abbott, Nate Sawtell, Manuel Beiran, and Kaushik Lakshminarasimhan for their comments on the manuscript. The M.J.W. laboratory is supported by the NINDS Intramural Research Program. A.L.-K. and S.M. were supported by the Gatsby Charitable Foundation, NSF award DBI-1707398, and the Simons Collaboration on the Global Brain. S.M. was also supported by the Swartz Foundation. A.L.-K. was also supported by the Burroughs Wellcome Foundation, the McKnight Endowment Fund, and NIH award R01EB029858.

## Methods

### 1 Network model

We model the input pathway to cerebellum-like structures as a three-layer feedforward neural network. The input layer activity ***x*** represents the task subspace (see below) and task-irrelevant activity. The representation ***x*** is sent to the compression layer via a compression matrix 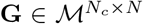. We consider both linear compression layer neurons, for which **c** = **G*x*** and ReLU neurons, for which **c** = [**G*x*** – ***θ***]_+_, where the rectification is applied element-wise. The output of the compression layer is sent to the expansion layer via a matrix 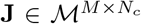, and we set ***m*** = *ϕ*(**Jc** – ***θ***), once again applied element-wise. In our result, *ϕ* is typically a Heaviside threshold function, except when considering nonlinear regression and when re-analyzing the data from [13], for which we used a ReLU nonlinearity. The nonzero entries of the expansion matrix **J** are sampled independently and i.i.d. from 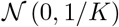, where *K* is the number of incoming connections onto an expansion layer neuron. The thresholds ***θ*** are chosen adaptively and independently for each neuron to obtain the desired coding level (fraction of active neurons) *f* or *f_c_* [6], for the expansion and compression layer respectively. The expansion representation ***m*** is read out via readout weights ***w***, i.e. *y^μ^* = ***w***^*T*^(***m*** – *f*) which are set using a Hebbian rule (Hebbian classifier), unless stated otherwise, i.e.

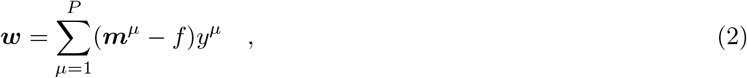

where *y^μ^* are the target labels and the subtraction is intended element-wise.

### Direct expansion network

To compare the performance of the bottleneck network to one without the compression layer, we also implement a *direct expansion* network, in which the input layer ***x*** is directly expanded to the expansion layer ***m*** via a sparse expansion matrix **J**, i.e. ***m*** = *ϕ*(**J*x*** – ***θ***). The matrix 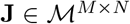 has *K* nonzero elements per row, with these entries sampled as for the bottleneck network.

## 2 Metrics of dimension and noise

To quantify the dimension of a representation, we use a measure based on its covariance structure [7, 42]. For a representation ***x*** with covariance matrix **C**^*x*^, which has eigenvalues λ_1_,…,λ_*n*_, we define

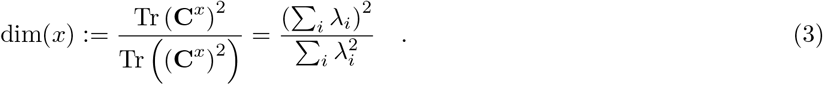

To quantify noise strength, we follow previous work and consider the Euclidean distance between a noiseless pattern 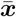 and a noisy one ***x***, specifically 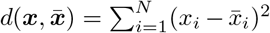 [6]. This distance is averaged over the noise distribution and normalized by the average distance among pairs of noiseless input patterns, leading to:

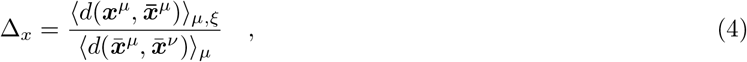

where *μ* denotes the average over the input distribution and *ξ* the average over the noise distribution. With this normalization, Δ_*x*_ = 1 if noisy patterns are on average as distant from their noiseless version as two different input patterns are with respect to each other. The definition of Δ_*c*_ and Δ_*m*_ for the noise strength at the compression and expansion layer is analogous to the one in Eq. 4.

## 3 Input representation of task-relevant variables

We model the input representation ***x*** as a linear mixture of task-relevant and task-irrelevant activity (i.e. noise). The task-relevant variables are described by a *D*-dimensional representation ***z***, and are encoded in the input layer via a matrix with orthonormal columns, i.e. 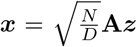 where 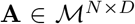. The columns of **A** can always be chosen to be orthonormal to each other, since we assume that *N* > *D*. The factor 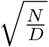 ensures that input layer activity is of order 1.

The task variables ***z*** consist of *D*-dimensional random Gaussian patterns, sampled from 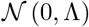, where Λ is a *D* × *D* diagonal matrix with diagonal elements λ_1_,…, λ_*D*_. The {λ_*i*_} represent the task subspace PCA eigenvalues, and to control their decay speed we set λ_*i*_ = *i*^−*p*^ and vary the parameter *p*. In the absence of noise, the input representation has rank *D*, and its covariance matrix **C**^*x*^ = **AΛA**^*T*^. Notice that because of the orthonormality of the columns of **A**, the nonzero eigenvalues of **C**^*x*^ are equal to λ_1_,…,λ_*D*_, i.e. the PCA eigenvalues of ***x*** are the same as ***z***. Therefore, in the absence of noise, dim(***x***) = dim(***z***). This can also be seen by noting that the columns of **A** form an orthonormal set, **A**^*T*^**A** = **I**, and using the cyclic permutation invariance of the trace, for any representation ***z*** with covariance matrix **C**^*z*^ (possibly non-diagonal):

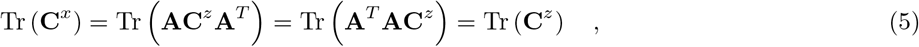

and analogously for Tr ((**C**^*x*^)^2^).

To model *distributed* input representations, we sample a random *N*-by-*N* orthogonal matrix **O** from a Haar measure (the analog of uniform measure for matrix groups), and select the first D columns of **O** to be the columns of **A**. To model a *clustered* input representation, we split the *N* input neurons into D groups. For simplicity, we take these groups to be equally sized, i.e. each group consists of *N_g_* neurons, but our results can be easily generalized to the case of groups with different sizes. The elements of the matrix **A** are set as

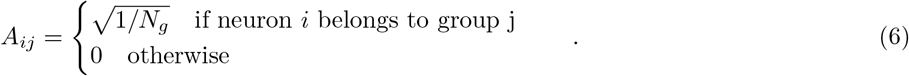

## 4 Noise model

We introduced two sources of noise in the model: input noise and expansion layer noise. For input noise, we considered mean-zero additive noise, i.e. we corrupt the noiseless input representation 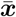 as

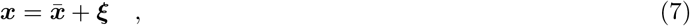

where ***ξ*** is a random vector with mean zero and covariance matrix **C**^*ξ*^. In contrast, the specific implementation of expansion layer noise depended on the type of nonlinearity used at the expansion layer. For binary neurons, used for most of our results, we randomly flipped a fraction *σ_m_* of the neurons for every pattern. For ReLU neurons, we added random isotropic Gaussian noise with variance 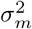 after the rectification, while keeping the final rate positive. Noise could also affect the compression layer representation, but we can absorb this contribution into the noise at the expansion layer (as noise at the expansion layer increases monotonically with the noise at the compression layer, see appendix B).

## 5 Random classification task

A random classification task is defined by first assigning binary labels *y^μ^* = ±1 at random to patterns 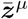, for *μ* = 1,…,*P*, in the task subspace. The network is required to learn these associations and generalize them to patterns which are corrupted by noise. When readout weights are learned using a Hebbian rule, 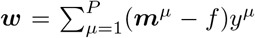, where the subtraction is intended element-wise, the probability of a classification error can be expressed in terms of the signal-to-noise ratio (SNR) of the input received by the readout neuron, as 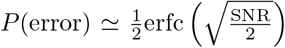 [6]. Previous work [7] has shown that the SNR can be expressed as

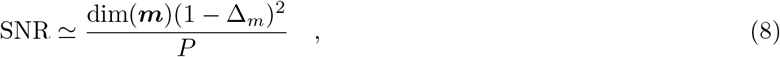

where Δ_*m*_ is the noise strength at the expansion layer, normalized by the average distance among noiseless patterns (see section B). On the other hand, dim(**m**) is the *noiseless* dimension, i.e. the dimension of the *task-relevant **m*** representation.

## 6 Compression architectures

Here, we briefly describe the different types of compression that we considered in the main text.

### 6.1 Random compression

We model random, unstructured compression by sampling the entries of the compression matrix **G** i.i.d. from a Gaussian distribution, i.e.

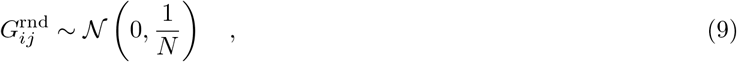

where the scaling of the variance is chosen to obtain order 1 activity in the compression layer.

### 6.2 Principal component-aligned compression

For principal component-aligned (PC-aligned) compression the rows of **G** are set equal to the task-relevant principal components of the input. Since the task-relevant variables are embedded in the input layer via the orthonormal columns of **A**, the latter are the task-relevant principal components. Therefore,

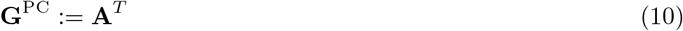

The above expression is valid when *N_c_* = *D*. If *N_c_* > *D*, we duplicate the rows of **G**^PC^, which results in a clustered representation at the compression layer. Note that with this type of compression, the *task-relevant* activity in the compression layer is *decorrelated,* because the task-relevant covariance matrix is given by **C**^*c*^ = **AC**^*x*^**A**^*T*^ = **C**^*z*^, which is diagonal by construction.

### 6.3 Whitening compression

To obtain a whitened spectrum, the rows of **G**^PC^ can be scaled in such a way that **C**^*c*^ = **I**. Since the eigenvalues of **C**^*x*^ are the same as the PCA eigenvalues of the task subspace representation ***z***, this is accomplished when:

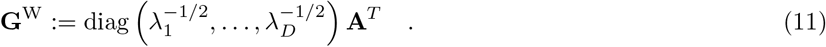

Similar to PC-aligned compression, if *N_c_* > *D*, we duplicate the rows of **G**^W^.

### 6.4 Optimal compression

We define the optimal compression matrix as either **G**^PC^ or **G**^W^, depending on which leads to the best performance. For nearly all the regimes we consider, whitening leads to the best performance.

### 6.5 Optimization of compression weights via gradient descent

For Fig. 2a,b, Fig. 4b, and Fig. S2c we used gradient descent to optimize the compression weights. We trained compression weights using backpropagation under the assumption that readout weights are learned using the Hebbian rule (Eq. 2). More precisely, for each epoch we sampled a random sparse expansion matrix **J** and *B* = 10 random classification tasks, each consisting of *P* = *D* = 50 target patterns.

For Fig. 2a,b and Fig. S2c, Hebbian readout weights are set independently for each task, after which the compression weights are updated in the direction that decreases the loss (binary cross-entropy) computed on noise-corrupted test patterns. The update step was performed using the Adam optimizer [43], with a learning rate *η* = 10^-4^. To facilitate learning by gradient descent, we replaced threshold nonlinearities in the expansion layer with ReLU nonlinearities. Adaptive thresholds were set, as in the rest of the paper, to obtain the desired coding level f. We used the same setup to test the performance in the presence of nonlinearities in the compression layer. We introduced ReLU nonlinearities in the compression layer in the same way as we did for the expansion layer, with a coding level *f_c_* = 0.3. The other parameters were *p* =1, *σ* = 0.1, *σ_M_* = 0.1, *f* = 0.1, *N* = 500, *N_c_* = 250, *M* = 1000 and *K* = 4.

For Fig. 4b, the compression weights were adjusted to maximize dimension and minimize noise at the compression layer, i.e. to maximize 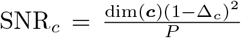. At every epoch, excitatory weights were constrained to be nonnegative. Isotropic Gaussian noise was added of strength *σ* = 0.2 was added to the input. Other parameters: *N_c_* = *D* = 10, *p* =1.

### 6.6 Hebbian compression

In Figs. 6, 7, we considered biologically-plausible learning of compression weights. In particular, we exploit the well known result that Hebbian plasticity leads to the post-synaptic neuron extracting the leading principal component of its input. In the presence of sparse compression connectivity, a compression layer neuron receives input from *L* input layer neurons. We call *S_i_* = {*j*_1_,…,*j_L_*} the set of indices that neuron *c_i_* receives input from. The covariance 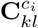 of such input is therefore a *L*-by*L* matrix given by

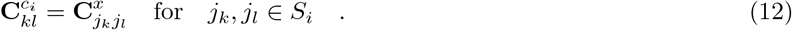

The leading principal component of the input to neuron *c_i_* is the (normalized) eigenvector of 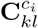 corresponding to its leading eigenvalues. Therefore, to mimic Hebbian plasticity in the presence of sparse connectivity we set the *i*^th^ row of **G** to the leading eigenvector of 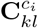. Notice that, for small *L*, the leading eigenvector of 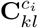 might be substantially different from the leading eigenvector of the full covariance of the input **C**^*x*^. Therefore, sparse connectivity introduces diversity of tuning across compression layer neurons, at the cost of pushing the tuning vectors outside of the task subspace, resulting in stronger noise.

## 7 Derivation of optimal compression

A key result of our theory is that, when expansion weights are random, optimal compression requires compression layer neurons to be tuned to task-relevant input principal components (PCs). Furthermore, the gain of different compression layer neurons could be adjusted to further increase performance. However, to what degree is convenient to do so depends on the input noise strength.

We start from the expression for the SNR of a Hebbian classifier (Eq. 8), which is a proxy of classification performance on a random classification task. To maximize the SNR, we would ideally maximize dim(***m***) while minimizing the noise Δ_*m*_. We find that aligning the weights to the PCs favors both objectives, while performing additional whitening increases dimension but also noise. While performance depends on dimension and noise *at the expansion layer*, in most cases the dimension and noise *at the compression layer* is sufficient to explain the resulting performance. This is because 1) noise at the expansion layer is a monotonic function of noise at the compression layer (see appendix B), and 2) dimension of the expansion layer depends on the dimension of the compression layer (see appendix A). However, dimension can also depend on the fine structure of the compression layer representation, in particular when the expansion connectivity is sparse (see appendix A). Below, we show how the properties of compression weights determine dimension and noise at the compression layer, motivating our definition of optimal compression.

### 7.1 Effect of compression on dimension

Here we present analytical results on the dimension of the compression layer in the case of linear compression. By definition of the dimension, we need to compute:

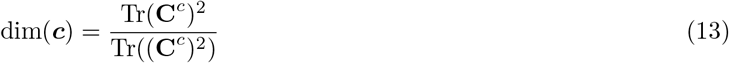

For random compression, it is convenient to reinterpret the trace as an average across compression layer neurons:

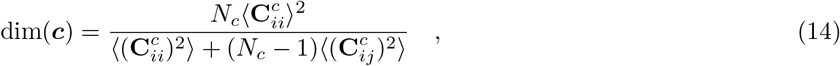

where the average is intended over *i* and *j*. We now define 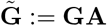, i.e. the effective matrix transforming the task variables into the compression layer representation (up to a factor 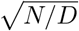, which is irrelevant for the dimension). Since the columns of **A** are orthonormal, the elements of 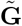 are also normally distributed and independent, with mean zero and variance 1/*N*. We have that 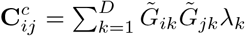. Approximating the average over *i* and *j* with the average over the distribution of 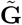, we obtain the dimension of ***c***:

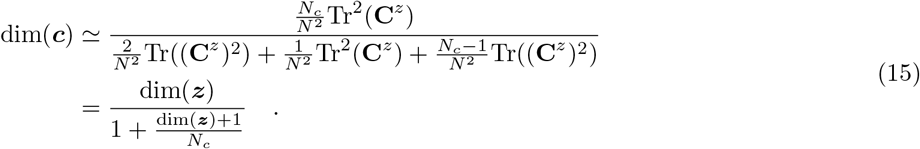

Notice that dim(***x***) = dim(***z***) since we assume an orthonormal embedding of the task subspace. Eq. 15 shows that random compression always reduces dimension, only preserving it in the limit of many compression neurons, *N_c_* ≫ dim(***x***). This is due to the distortion of the input layer representation introduced by the random compression weights [6] (Fig. 2e).

However, such distortion can be avoided by a choice of compression matrix that preserves the geometry of the input representation and its dimension (dim(***c***) = dim(***x***)). These compression matrices are characterized by orthonormal rows, a more stringent requirement that cannot be guaranteed by each compression layer neuron sampling its inputs independently. To see this, consider the traces appearing in the dimension expression (Eq. 13), which can be written using the effective matrix 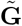 introduced above:

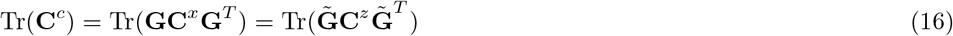

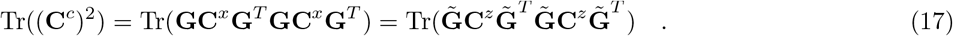

Thanks to the cyclic permutation invariance of the trace, computations of the traces in Eqs. 16, 17 above boils down to the computation of 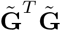. In particular, if 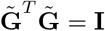, the traces will be unaffected by 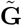 and dim(***c***) = dim(***z***). One situation in which this happens is when **G** satisfies two conditions: 1) the rows of **G** are orthonormal and 2) the columns of **A** are contained in the span of the rows of **G** (see appendix D for the proof). We call such compression “orthonormal” compression. Intuitively, the orthogonality of the rows of **G** avoids any distortion of the representation, while the columns of **A** need to be in the span of the rows of **G** to avoid that part of the task subspace being filtered out during compression.

PC-aligned compression is a special case of orthonormal compression, in which the rows of **G** are aligned with the columns of **A**. If the expansion connectivity is dense, i.e. **J** is a fully-connected matrix, such alignment will not lead to any improvement compared to any other orthonormal compression. However, when the expansion connectivity is very sparse, PC-aligned compression leads to larger dimension at the expansion layer (Fig. 3e). Since expansion connectivity in cerebellum-like structures is very sparse, we included the alignment with PCs as a feature of optimal compression. In general, studying the dimension of the expansion layer representation in the sparse connectivity scenario analytically is challenging, and we therefore study it either numerically or using semi-analytical Monte Carlo integration.

Finally, compression can also increase dimension. This can occur for whitening compression, which equalizes the variances of different input principal components. Because we consider linear compression, the task-relevant dimension of the compression layer is bounded by *D*. This bound is attained by whitening compression, as by definition it results in **C**^*c*^ = **I**. In summary, the effect of compression on dimension can be: 1) beneficial, if dim(***c***) > dim(***x***), e.g. for whitening compression 2) neutral, if dim(***c***) = dim(***x***), i.e. for orthonormal compression or 3) detrimental, if dim(***c***) < dim(***x***), e.g. for random compression.

### 7.2 Effect of compression on noise

Input noise can be separated in *task noise*, which corrupts the input representation along the task subspace; and *task-orthogonal noise*, which lies in directions orthogonal to the task subspace. Compression cannot attenuate task noise without reducing the signal strength as well, nor can it attenuate expansion layer noise. However, it can filter out noise in task-orthogonal directions. The extent to which this happens depends on the alignment between the incoming weights onto compression and the task subspace, as we show below.

We start by computing the noise strength at the input layer. From Eq. 4, we see that to obtain the noise strength we need to compute 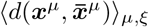, and 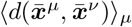. For this noise model, 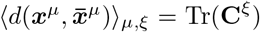. To compute the average distance among noiseless patterns, we notice that

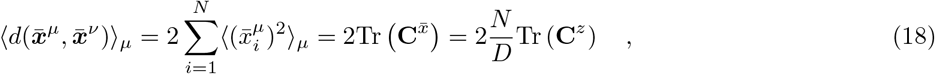

where 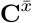 is the trace of the covariance matrix of the *noiseless* input. The noise strength can therefore be written as

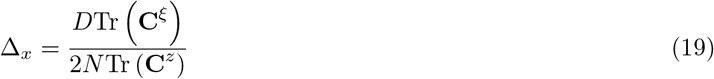

We now use the same technique to compute the noise strength at the compression layer. By direct calculation,

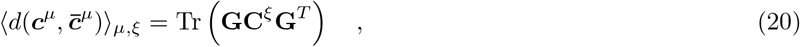

i.e. the average distance between noiseless patterns and their noisy realizations depends on the alignment between the rows of **G** and the directions along which noise varies (the eigenvectors of **C**^*ξ*^). Similarly,

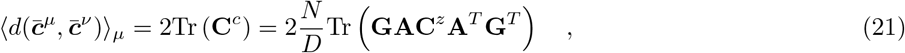

i.e. the average distance among noiseless patterns depends on the alignment between the rows of **G** and the columns of **A** (which define the task subspace). Notice that the factor 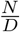 results from the fact that we want order 1 input layer activity (see section 3).

In most of the applications, we consider the case of isotropic noise, i.e. **C**^*ξ*^ = *σ*^2^**I**. In this scenario, we expect the noise component along the task manifold to scale as *D/N*. If the noise strength at the input layer is Δ_*x*_, the minimum noise level achievable without signal reduction is therefore given by

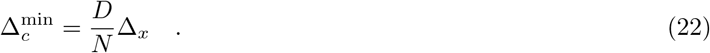

This means that for fixed task manifold representation, 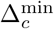 scales as 1/*N*, i.e. the more redundant the input representation is, the more it is possible to denoise it. PC-aligned compression attains this minimum noise strength. Indeed, since **G**^PC^ = **A**^*T*^,

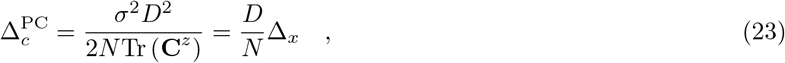

where we used Eq. 19 in the case of isotropic noise for the second equality.

In contrast, if compression weights are chosen randomly, the compression matrix rows are equally likely to overlap with task subspace and task-orthogonal directions, and the noise strength remains unchanged on average: 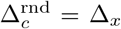. Indeed, for random compression one has that, when averaging over the weights, 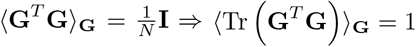, and 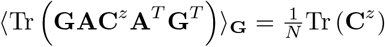. We can therefore write that, on average,

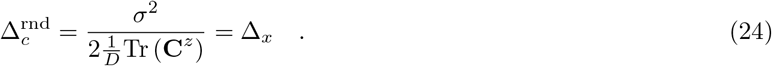

We have seen that whitening compression leads to the largest increase in dimension. However, by increasing the variance of sub-leading components to achieve normalization, whitening might also inflate the effect of noise. Indeed, for whitening compression we set 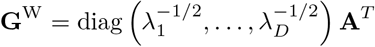, for which we get 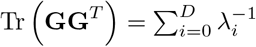 and, for the denominator, Tr (**C**^*c*^) = Tr(**I**) = *D*. The resulting noise strength is therefore given by

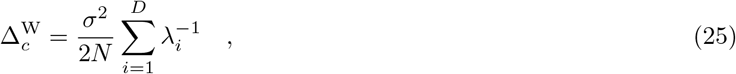

where *σ*^2^ is the variance of the input noise. When **C**^*z*^ has a decaying eigenvalue spectrum, 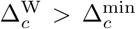. This detrimental effect of whitening is particularly strong if the signal eigenvalues decay very quickly or if the noise is very strong.

In summary, we showed that random compression leaves isotropic noise unaffected, PC-aligned compression attains the maximum noise reduction, and whitening compression, while filtering out noise in directions orthogonal to the task subspace, might inflate noise along the task subspace directions.

## 8 Optimal compression in the insect olfactory system

### Recurrent compression layer

We model recurrent interactions in the compression layer via the differential equation

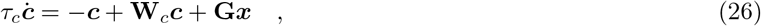

where **W**_*c*_ is the matrix of recurrent interactions in the compression layer and **G** is the matrix of feedforward interactions from the input to the compression layer. We assume that *τ_c_* is much smaller than the timescale at which the input varies, so that we can focus on the steady state dynamics given by

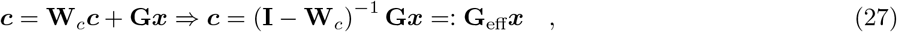

where we defined the effective feedforward matrix as **G**_eff_ := (**I** – **W**_*c*_)^-1^ **G**. Therefore the compression matrix **G**_eff_ can be thought as the effective steady state compression matrix in the presence of recurrent interactions and linear neurons. This factorization can be used to interpret optimal compression weights as a combination of feedforward and recurrent interactions. For the antennal lobe,

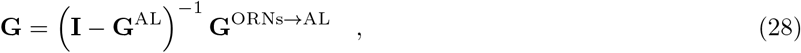

where **G**^ORNS→Al^ describes the projections from the antenna to the antennal lobe while **G**^AL^ captures the interglomeruli interactions.

### Recurrent inhibition

Here, we study the biologically relevant case of purely inhibitory recurrent connectivity. We start by considering global inhibition, characterized by a rank-one connectivity matrix that we write as

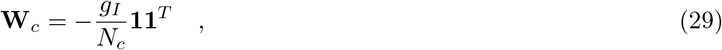

where **1** is the vector of all ones. In this case, the inverse of **I** – **W**_*c*_ can be computed explicitly using the Sherman–Morrison formula:

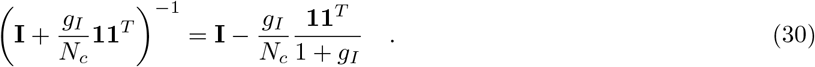

Plugging this expression in the definition of **C**^*c*^,

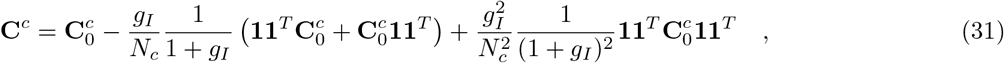

where 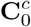 is the covariance matrix of c without considering recurrent inhibition. We now define ***u***^1^ := **U**^*T*^**1**, i.e. the vector of the projections of the eigenvectors of 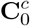 on the constant mode **1**. With this definition,

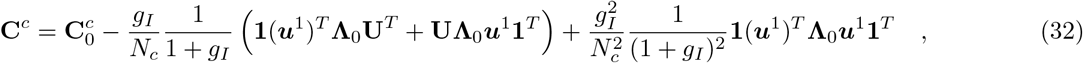

where **Λ**_0_ is a diagonal matrix containing the eigenvalues of the 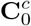. If **1** is one of the eigenvectors of 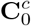, then **u**^1^ only has one nonzero entry. In this case, global inhibition controls the strength of the uniform mode in **C**^*c*^, and can be used to set it to zero. More generally, Eq. 32 shows that global inhibition acts on the projection of input modes on the constant mode. This is equivalent to say that the effect of global inhibition on a certain mode depends on the mode mean, i.e. the average of the eigenvector entries. It is straightforward to generalize this derivation to the case of multiple inhibitory neurons which act on non-overlapping groups of neurons.

### Realistic odor sensory neuron responses

To generate realistic responses of olfactory sensory neurons (OSNs), we considered a widely used dataset containing the responses (difference from baseline firing rate) of 24 types of OSNs to a panel of over 100 odors [21]. We use these responses to estimate the covariance across different odor receptor types, thereby obtaining an estimate of the values of different blocks in the covariance matrix of the input layer **C**^*x*^ = **C**^OSN^ (Fig. S3a). While this estimate of the covariance matrix is noisy due to the limited number of odors in the dataset, we nonetheless found that the off-diagonal elements of **C**^OSN^ were on average more positive than expected by chance given the amount of noise in the estimate, by shuffling the responses of OSNs to different odors (Fig. S3b).

In Fig. 3c, d, we set **G** to mimic the convergence of OSNs of the same type:

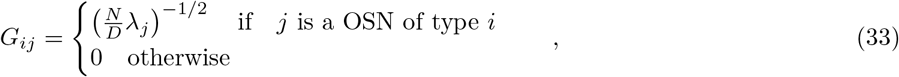

where we divided by λ_*j*_, the diagonal elements of **C**^OSN^, to implement normalization mechanisms across different types of inputs. To implement global inhibition, we set 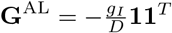 in Eq. 28. The dimension of the compression layer representation increases monotonically with *g_I_*, but saturates around *g_I_* ~ 1. We therefore set *g_I_* = 10 to ensure the strongest effect of global inhibition. For Fig. 3d, we modeled responses to mixture of odors as random patterns sampled from a Gaussian distribution with mean zero and covariance matrix **C**^OSN^. Therefore, we sampled 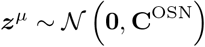 for *μ* = 1,…,*P*, embedded these patterns in the input layer using a clustered embedding, and use them as the training set for a random classification task. For testing, we added Gaussian isotropic noise to the input layer **C**^OSN^ with standard deviation *σ_x_* = 0.5.

## 9 Local decorrelation

In Fig. 5, we considered the scenario in which expansion layer neurons in the direct expansion architecture could *locally decorrelate* their input (i.e. perform whitening of their inputs), and nonlinearly mix the resulting signals. Using the same argument as for Hebbian compression with sparse connectivity, the covariance of the input to an expansion layer neuron *m_i_* is

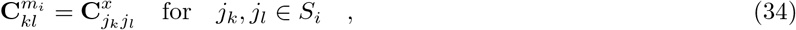

where *S_i_* is the set of *K* afferents to neuron *m_i_*. We implemented local decorrelation by assuming that the *K* weights of the incoming connections on an expansion layer neuron are set as

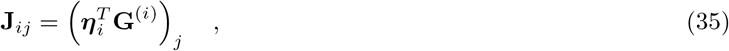

where ***η***_*i*_ is a *K*-dimensional Gaussian random vector with independent entries and **G** ^(*i*)^ is such that 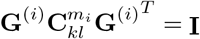. In words, each expansion layer neurons performs whitening of its input via the matrix **G**^(*i*)^ and mixes the resulting inputs with random coefficients *η_ij_*.

### 9.1 Supervisory input from DCN to the pontine nuclei

To model supervisory input from DCN affecting plasticity at cortico-pontine synapses, we assume that the DCN output is close to the target output, i.e. DCN(*t*) ≃ *y*(*t*). We consider a modification of Oja’s rule:

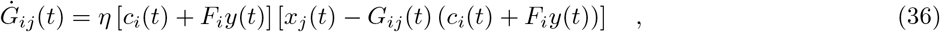

where *F_i_* is a constant determining the strength of the supervisory input. In simulations, we sample *F_i_* randomly and independently for each compression layer neuron. This form of plasticity can be interpreted as adding DCN input to the compression layer representation *only for plasticity purposes,* without changing the network dynamics. For simplicity, consider the effect on a single compression layer neuron *c* := *c_i_*, assuming that the target is a scalar, i.e. *y* ∈ ℝ. Assuming that the weight evolution is slow, so that we can average over ***x***,

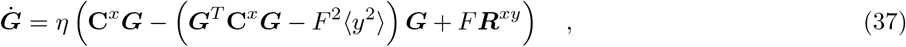

where ***R***^*xy*^ := 〈***xy***〉_*x*_ is the vector of input output correlations, with one component for each input dimension, and ***G*** denotes the vector of incoming weights onto *c*. We can express both ***G*** and ***R***^*xy*^ in the basis formed by the PCA eigenvector of ***x***, i.e. 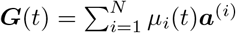 and 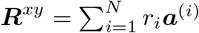. We can then rewrite Eq. 37 as

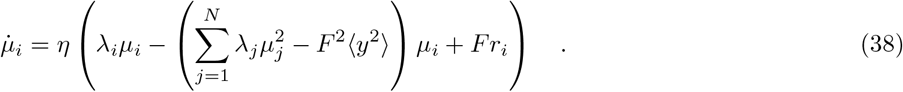

This equation shows that the overlap *μ_i_* of the weight vector with a certain input PC ***a***^(*i*)^ will be driven by how correlated that PC is with the target *y* (last term on the RHS). Therefore, if *F* is large enough, even leading PCs which are uncorrelated with the target will not be extracted by compression layer neurons.

### 10 Forward model learning

When learning a forward model, the network should learn to predict the sensory consequences of motor commands. We assume that the dynamics of a motor plant can be summarized by a set of differential equations

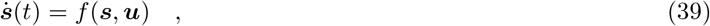

where ***s*** ∈ ℝ^*N_s_*^ describes the sensory state associated with the plant (such as proprioceptive or visual feedback), ***u*** ∈ ℝ^*N_u_*^ is the motor command, and *f* is a smooth, vector-valued nonlinear function that summarizes the dynamics of the plant. The forward model task is then to predict ***s***(*t* + Δ), given ***s***(*t*) and ***u***(*t*). If the time interval Δ is small to compared to the speed of the plant dynamics, we can approximate

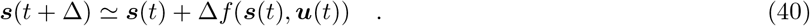

We assume that the cerebral cortex sends to the cerebellum information about both ***s***(*t*) and ***u***(*t*). To implement a forward model, the cerebellum should relay the information received by the cortex (first term in Eq. 40), and add to it the nonlinear function *f* (***s***, ***u***). We assume that the relay operation is carried out by the mossy fiber to DCN pathway, while the Purkinje cells compute the negative of the nonlinear term and feed it to the DCN. The target of learning at Purkinje cells is then given by ***y***(*t*) = – *f*(***s***(*t*), ***u***(*t*)).

In the model, we concatenate ***s***(*t*) and ***u***(*t*) in a single vector of task variables ***z***(*t*) ∈ ℝ^*N_s_*+*N_u_*^ and embed it in the input representation in a distributed fashion. The target ***y*** of forward model learning is a vector, with an entry for each degree of freedom of the motor plant. In simulations, we only consider one target entry at the time, i.e. we assume that different target components are learned by separate sets of Purkinje cells, and report the average performance.

In summary, we cast the forward model learning task into a nonlinear regression task, in which the network has to learn a nonlinear target function *f*(***z***) of the input ***z***. Because the target function is smooth, it is convenient to use ReLU neurons the compression layer instead of binary neurons. Furthermore, since Hebbian learning of the readout weights performs poorly in regression tasks, we used a pseudoinverse learning rule, which is analogous to performing ridge regression

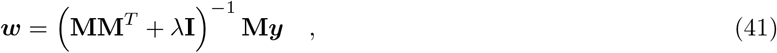

where λ is the ridge regularization parameter.

#### 10.1 Random target functions from Gaussian process priors

In Fig. S7, we considered an ensemble of target functions sampled from a Gaussian process (GP) prior. We set the mean function of the GP to be zero and its covariance function to be given by

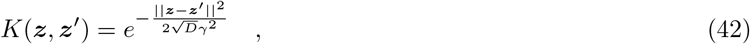

where *γ* is a parameter that controls how quickly-varying the sampled functions are.

#### 10.2 Planar two-link arm target

We consider dynamics of a two-joint arm in the absence of gravity (planar) [44]. The arm consists of two bars of length *l* and mass *m*, and its state is defined by the two joint angles *θ*_1_ and *θ*_2_, and by the corresponding angular velocities. The dynamics equations are written, in matrix form, as

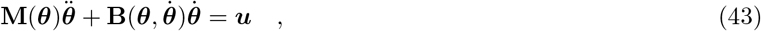

where ***u*** contains the two torques and **M** is a two-by-two matrix that contains the inertial terms

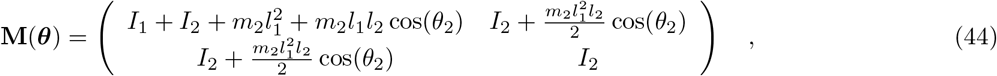

where *I*_1_ and *I*_2_ are the moments of inertia of the two links. The matrix **B** is given by

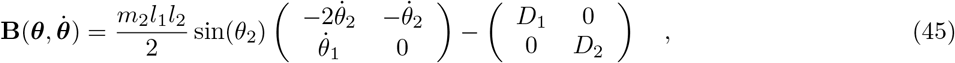

where *D*_1_ and *D*_2_ control the damping strength.

### 11 Analysis of simultaneous cortico-cerebellar recordings

#### 11.1 Summary of experimental setup

The recordings analyzed in Fig. 7 were performed using the experimental setup described in detail in [13]. In brief, *Ai93/ztTA/Math1-Cre/Rbp4-Cre* quadruple transgenic mice were head-fixed and performed pushed a handle to perform L-shaped trajectories, either to the left or to the right. We only considered data from the first session after a mouse was considered expert on the task. Furthermore, we only retained “pure” turn trials, i.e. trials in which mice did not push the handle in the incorrect lateral direction by more than 500 *μ*m at any point during either the forward or lateral motion segments. During the task, neural activity from layer-5 pyramidal neurons in premotor cortex and cerebellar granule cells was monitored simultaneously using two-photon microscopy, with a 30Hz sampling rate. More precisely, granule cells were imaged through a cranial window on top of lobules VI, simplex, and crus I.

#### 11.2 Model and input representation

To estimate task-relevant and task-irrelevant activity from the cortical recordings, we regressed the cortical activity using a set of basis functions aligned to the turn point in the behavioral trajectories. More precisely, we used two boxcar functions covering at most one second before the turn and two boxcar functions covering at most one second after the turn. Furthermore, we used separate basis functions for right and left turns. In total we then used eight boxcar basis functions, whose length was adapted to each trial trajectory. We considered task-relevant the activity that could be predicted using a linear model with such basis functions as predictors. All the residual (unpredicted) activity was deemed task-irrelevant.

The unobserved population followed the same update equations as the observed one. However, instead of having the measured cortical activity as the input, we generated synthetic data based on the measured task-relevant and task-irrelevant statistics. Synthetic task-relevant activity was generated using the linear regression model described above as a generative model. To sample task-irrelevant activity, we measured the sample covariance matrix of task-irrelevant activity separately for each session, and use it to generate new task-irrelevant activity for the unobserved population, assuming Gaussian statistics. Importantly, task-relevant and task-irrelevant activity was respectively weighted by two parameters, 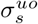 and 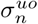.

#### 11.3 Measures of correlation and selectivity

The correlations in Fig. 7c were measured by computing the Pearson correlation coefficient among all neuron pairs. These correlation coefficient had mean zero across neuron pairs, therefore we took their standard deviation across neuron pairs as the measure of correlation strength for a single session, and then averaged across sessions. The same procedure was applied to correlations among granule cells and between granule cells and layer-5 cells, both in data and in the random and Hebbian model.

Our measure of cell selectivity to left/right turns is analogous to the one used in [13]. In particular, we devise an encoding model using four boxcar basis functions corresponding to before/after left/right turns. Each boxcar function was at most 300 ms long. After fitting the linear regression model with these basis functions, we quantified the number of coefficient significantly different from zero, independently for each of the 4 basis functions (criterion: *p* < 0.01), and normalized it by the number of neurons.

## A Dimension of the expansion layer

We define ***h*** := **J*c*** i.e. the input currents to the expansion layer neurons. The ***h*** representation has covariance matrix **C**^*h*^. However, to compute the dimension of the expansion layer it is not sufficient to compute the dimension of ***h***, because we need to take into account the nonlinearity of expansion layer responses. Analogously to Eq. 14, we can express the dimension of ***m*** as

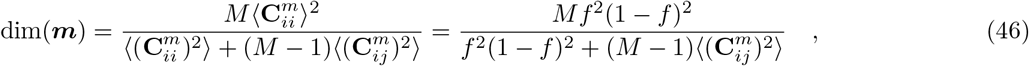

where the second equality follows from the expansion layer having a fixed coding level. Eq. 46 highlights that dimension is a function of the distribution of pairwise covariance among neurons. For fixed coding level *f* and threshold nonlinearities the covariance between two expansion layer neurons 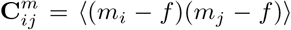 depends only on *f* and on the Pearson’s correlation between the respective input currents 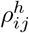:

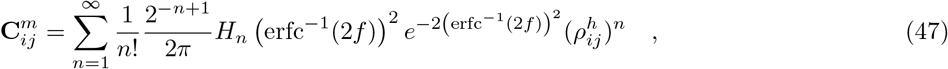

where 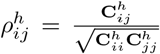, erfc^−1^ is the inverse of the complementary error function and *H_n_* is the *n*-th (physicist) Hermite polynomial. Therefore the problem of computing 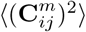 can be rephrased, for threshold nonlinearities, to the problem of computing all the moments of 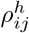. For typical regimes encountered in this paper, a truncation of Eq. 47 to the first few terms provides a good approximation of 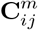, providing a much more efficient way to compute 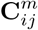 compared to its integral formulation (see for example [7], Eq. 43).

To compute 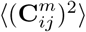, we need to estimate the moments of 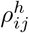 (across neurons). If all the weights in the network are sampled from mean-zero distributions, 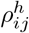 will also have mean zero and its distribution will be symmetric around zero. As a result, we only need to compute the even moments of 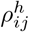. When other approaches are not feasible, one can directly estimate the required moments of 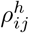 (typically the second and fourth moments are sufficient) from their empirical distribution for a particular network realization. In some cases, however, it is possible to obtain good approximations of 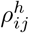 by considering the Gaussian approximation of the joint distribution of 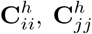 and 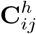, as we discuss below.

### 1 Fully-connected random expansion

For a fully connected random expansion with 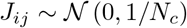, we find that the joint distribution of the elements of the input current covariance matrix, can be approximated by

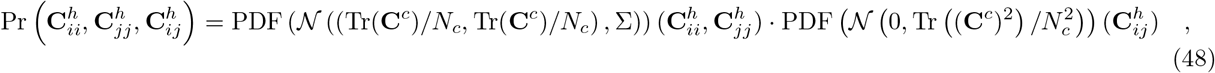

i.e. the probability density factorizes in a two-dimensional normal distribution in 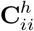 and 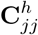 and a one-dimensional normal distribution in 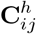. The covariance matrix of the two-dimensional Gaussian is given by

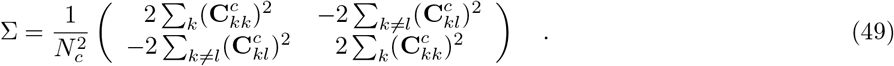

To find the first two terms in the expansion of 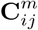, we will find approximations of 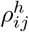 which are valid in the large *N_c_* regime, as long as dim(***c***) is not too small. Since 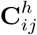 is independent of the diagonal elements, we write

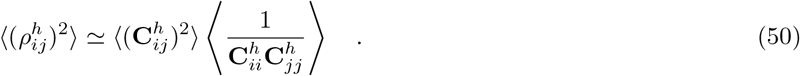

We now expand the reciprocal above up to second order in dim(***c***), and get

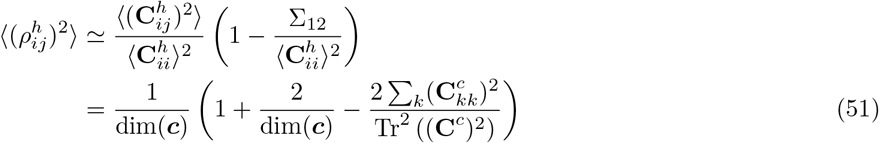

One can perform a similar calculation to compute 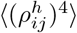, and find that, at the same order,

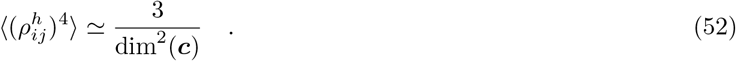

Using these results and the first two coefficients of the series in Eq. 47, one finds

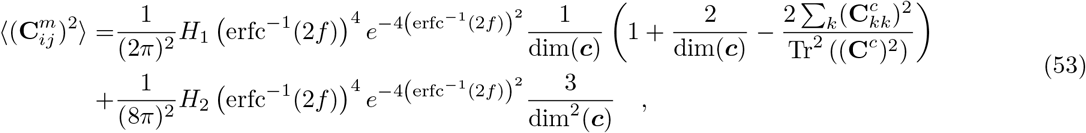

from which one easily obtains the dimension. Notice that, at first order, 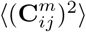 depends only on dim(***c***) and as a result dim(***m***) scales approximately linearly with dim(***c***). Even though the above is an approximation, we find empirically that in most cases dim(***c***) is one of the strongest factors affecting dim(***m***).

### 2 Sparsely-connected random expansion

When considering a sparsely connected expansion, the probability density in Eq. 48 has the same form, but the statistics instead of depending on the full matrix **C**^*c*^ depend on the input covariance matrix “seen” by neurons *i* and *j* of the expansion layer, i.e. on **C**^*c*^ sub-sampled to the pre-synaptic partners of neurons *i* and *j*. To obtain an approximation of 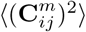, one has to compute the averages numerically in a Monte Carlo fashion. We find that for very sparse connectivity the degree at which the compression weights are aligned with PCs strongly affects the dimension of the expansion layer (Fig. 3e).

## B Noise at the expansion layer

Here we show that the noise strength at the expansion layer is a monotonic function of noise at the compression layer (excluding additional noise sources at the expansion layer). Following previous work [6], the noise strength at the expansion layer Δ_*m*_ can be written, for binary expansion layer neurons with fixed coding level *f*, as

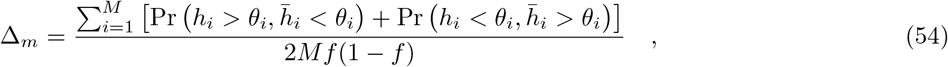

where *h* and 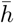 are the noisy and noiseless input current to the expansion layer neuron *i*, respectively. We treat *h_i_* and 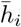 as correlated Gaussian variables with mean zero, variance *σ_i_* and 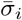 respectively and correlation coefficient *ρ_i_*. We have that

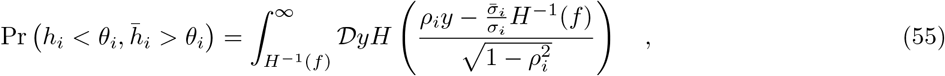

where 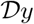 indicates the standard Gaussian measure and *H*(*f*) is the Gaussian tail probability, i.e. 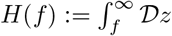. Because of symmetry, we have that 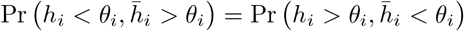, and

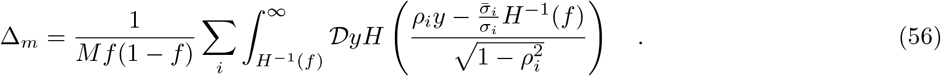

Under the approximation that all the expansion neurons are statistically equivalent, the sum over *i* can be simplified with the factor *M* at the denominator. Furthermore, the correlation coefficient *ρ* can be approximated by

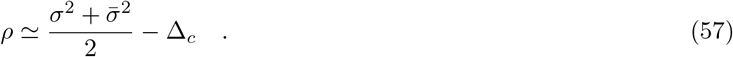

## C Using recurrent weights to adjust the input eigenvalue spectrum

We call 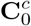 the covariance matrix of the feedforward input, i.e. 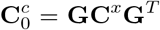. Since 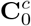 is a covariance matrix, it is diagonalized by a set of orthonormal eigenvectors, i.e. 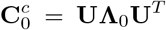, where the columns of **U** are the eigenvectors. Suppose we want to find **W**_*c*_ such that the covariance of the compressed representation has some desired eigenvalue spectrum 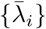. If we do not aim to change the eigenvectors, we can write the desired covariance matrix as 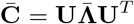 and make the ansatz 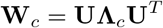, so that we need to solve

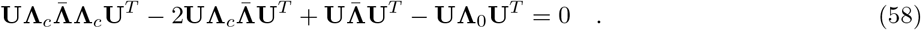

Since all the matrices share the same eigenvectors, we need to only solve the following equation

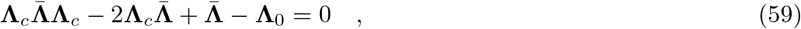

which can be solved for each eigenvalue of **W**_*c*_ separately, yielding

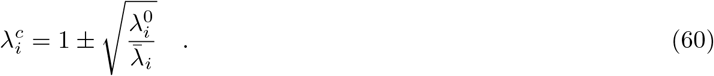

Notice that of these two solutions only 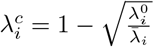 corresponds to a stable fixed point.

## D Preservation of dimension by orthonormal compression

### Proposition.

If the rows of **G** are orthonormal and ***a***_*i*_ ∈ span (***G***_1_,…, ***G***_N_*c*__), then the columns of 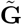 are orthonormal.

### Proof.

We can express the columns of **A** as 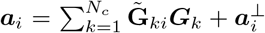. Since the rows of **G** are orthonormal, the orthonormality condition of the columns of **A** becomes:

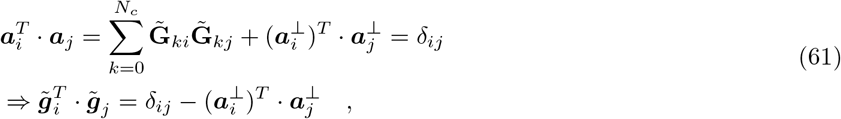

Since ***a***_*i*_ ∈ span(***G***_1_,…, ***G****_N_c__*), for all *i* = 1,…, *D*, then 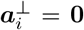 for all *i* = 1,…, *D*, and the columns of 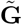 are orthonormal. This concludes the proof.

While not relevant for the proof above, it is worth noticing that the rows of 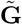 are also not orthonormal in general. We separate the rows ***G***_*i*_ ∈ ℝ^*N*^ of **G** in their component that lies in the task subspace and the one that is orthogonal to it

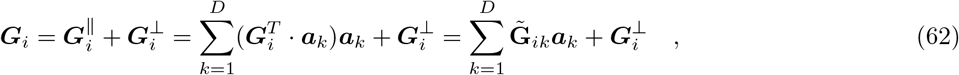

where ***a***_*k*_ is the *k*-th column of the matrix **A** and the last equality follows from the definition of 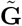. Since the rows of **G** are orthonormal, then we have that

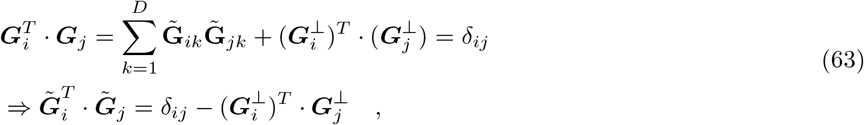

from which we conclude that the rows of 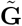 are also not orthonormal in general.

## Supplemental figures

**Figure S1:**
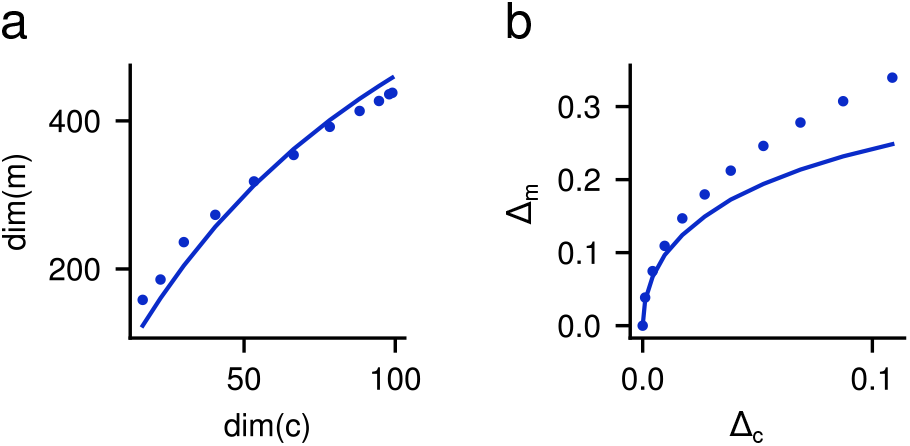
Expansion layer dimension and noise strength depend on compression layer dimension and noise strength. **a:** Dimension of the expansion layer representation as a function of the compression layer one. The compression layer representation was distributed, and its dimension was varied by changing p between 0 and 1. **b:** Noise strength Δ_*m*_ at the expansion layer as a function of the noise strength at the compression layer. Noise was additive, Gaussian, and isotropic at the compression layer, with standard deviation varying from 0 to 0.1. In both panels, solid lines show the theoretical result and dots are simulation results. Parameters: Nc = 100, *M* = 1000, *f* = 0.1.

**Figure S2:**
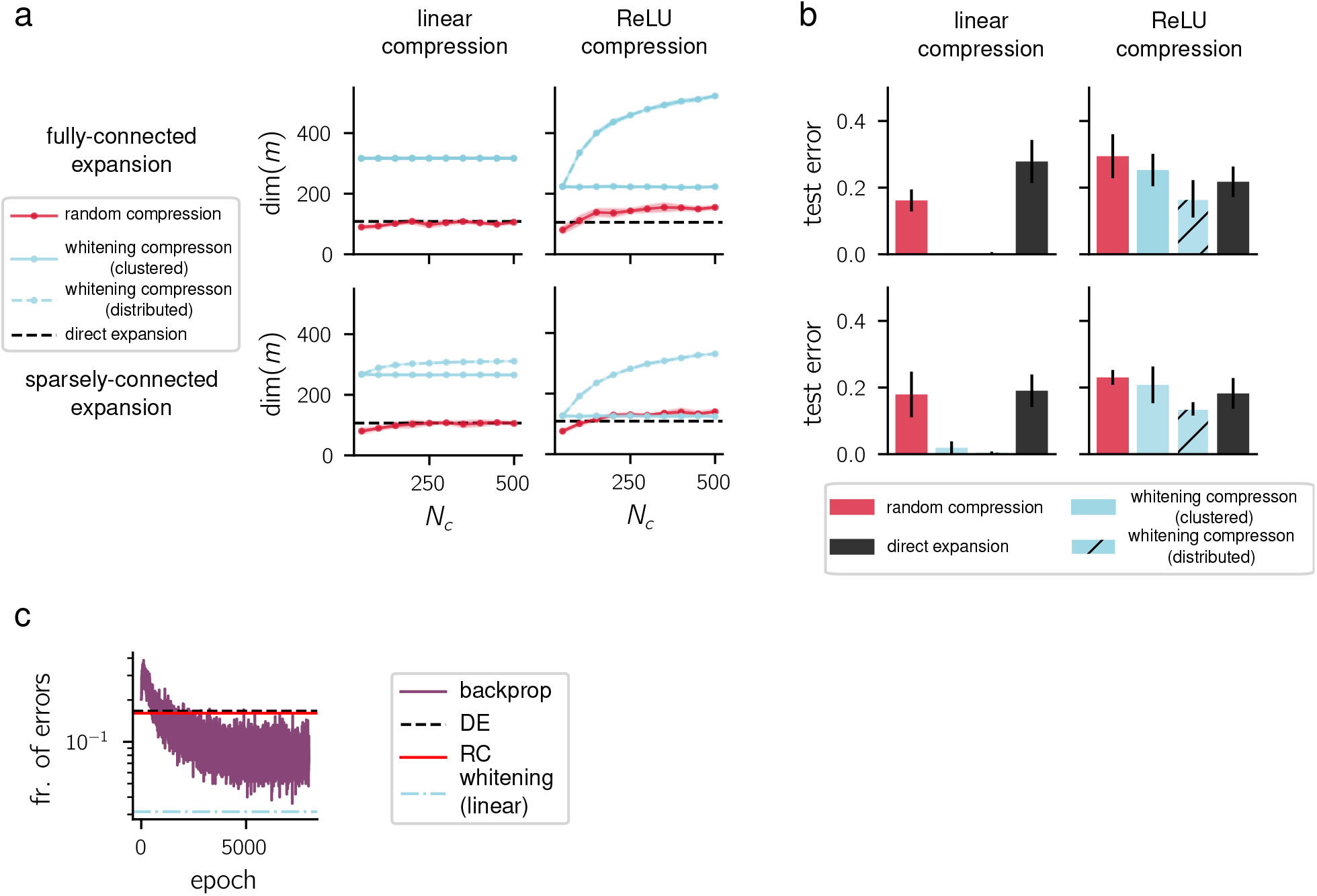
Effect of nonlinearities at the compression layer. **a:** Dimensionality of the expansion layer representation for fully-connected versus sparsely connected expansion (first versus second row) and for linear versus nonlinear (ReLU) compression. Shading indicates mean ± standard deviation. **b:** Same as **a**, but showing the generalization error in random classification. Errorbars indicate standard deviation. For both panels, the coding level of the compression layer was set to *f_c_* = 0.3 and the other parameters were *p* = 1, *D* = 50. **c:** Classification error as a function of training epoch when compression weights are trained using error backpropagation. Trained, direct expansion and random compression architectures all have ReLU nonlinearities. For comparison, we also plot the performance of whitening with a linear compression layer.

**Figure S3:**
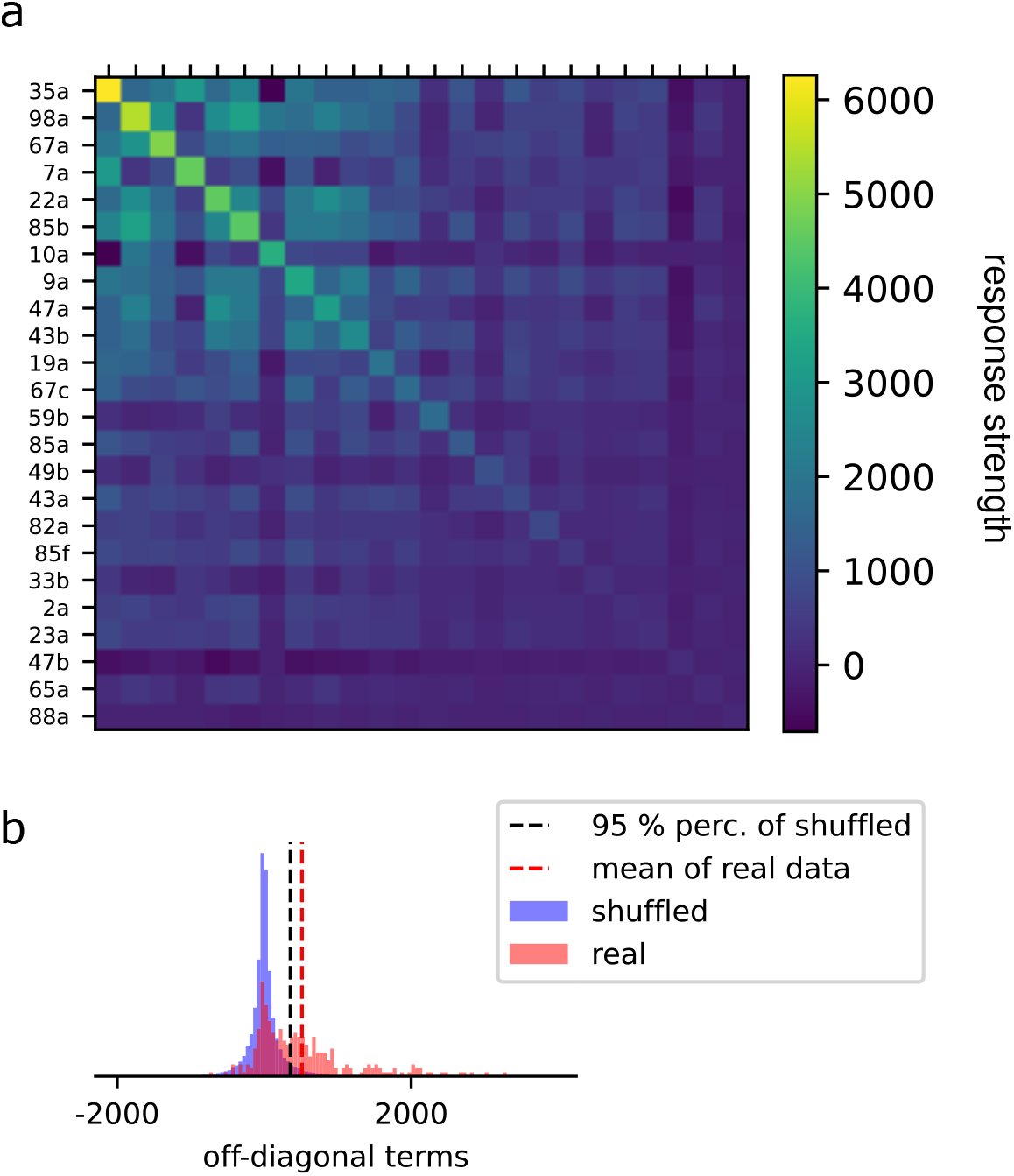
Realstic properties of odor receptor responses. **a:** Covariance of single odor receptors responses, computed from the Hallem-Carlson dataset [21], sorted according to the response variances. **b:** Histogram of off-diagonal terms in the covariance matrix in **a** (in red), compared to a shuffle distribution (blue) obtained by shuffling the responses to different odorant for a given odor receptor.

**Figure S4:**
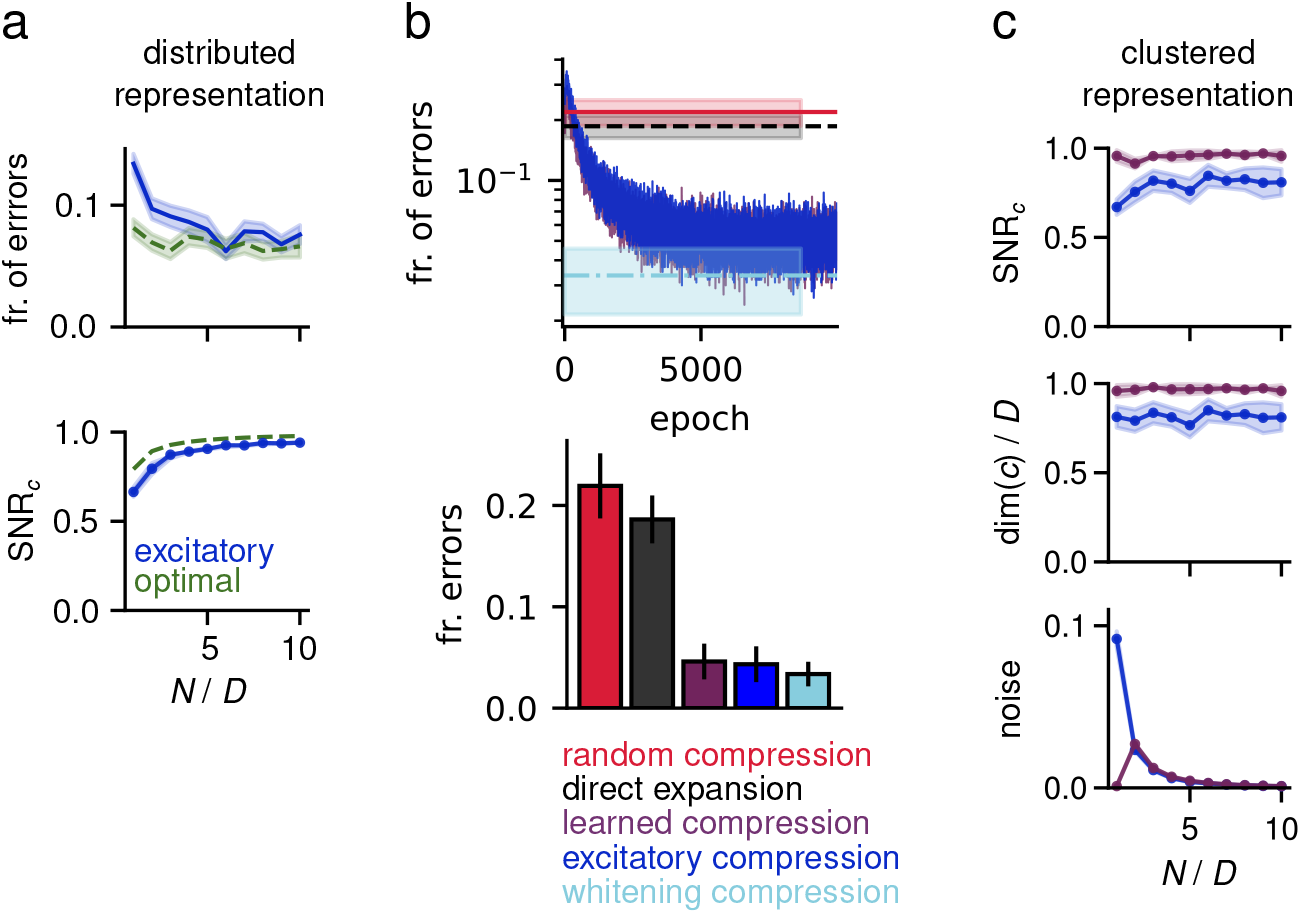
Sign-constrained compression for clustered and distributed representations. **a:** Fraction of errors on a random classification task (top) and signal-to-noise ratio of a Hebbian classifier reading out from the compression layer representation, i.e. SNR_*c*_ = dim(*c*)(1 – Δ_*c*_)^2^. Parameters are the same as in Fig. 4b. **b:** Results of gradient descent training of sign-constrained compression, when the loss function is given by the binary cross entropy of a Hebbian classifier reading out from the expansion layer representation. The procedure is the same used in Fig. 2a, b. **c:** Recurrent inhibition at the compression layer is necessary when the input representation is clustered. Results of gradient descent training with loss function given by SNR_*c*_, when the input representation is clustered with non-zero off-diagonal entries.

**Figure S5:**
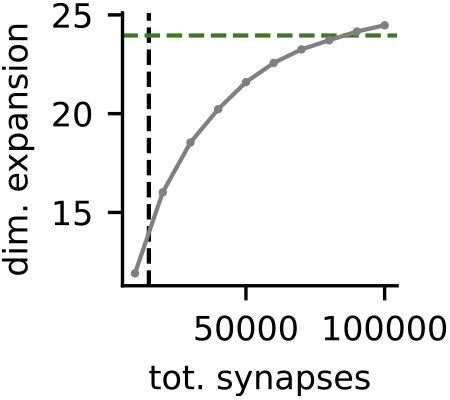
Optimal compression outperforms local decorrelation in direct expansion architecture for antennal lobe input and architecture. Dimesionality expansion dim(*m*)/dim(*x*) as a function of the total number of synapses, for the *direct expansion* architecture with local decorrelation at the expansion layer, as in Fig. 5. The green dashed line indicate the value obtained with optimal compression, which uses a total number of synapses indicated by the black vertical dashed line. The parameters were chosen to be consistent with the insect olfactory system anatomy, i.e. *D* = *N_c_* = 50, *N* = 1000, *M* = 2000, *p* =1, *f* = 0.1. To change the wiring cost of the local decorrelation strategy, we varied *K* between 5 and 50.

**Figure S6:**
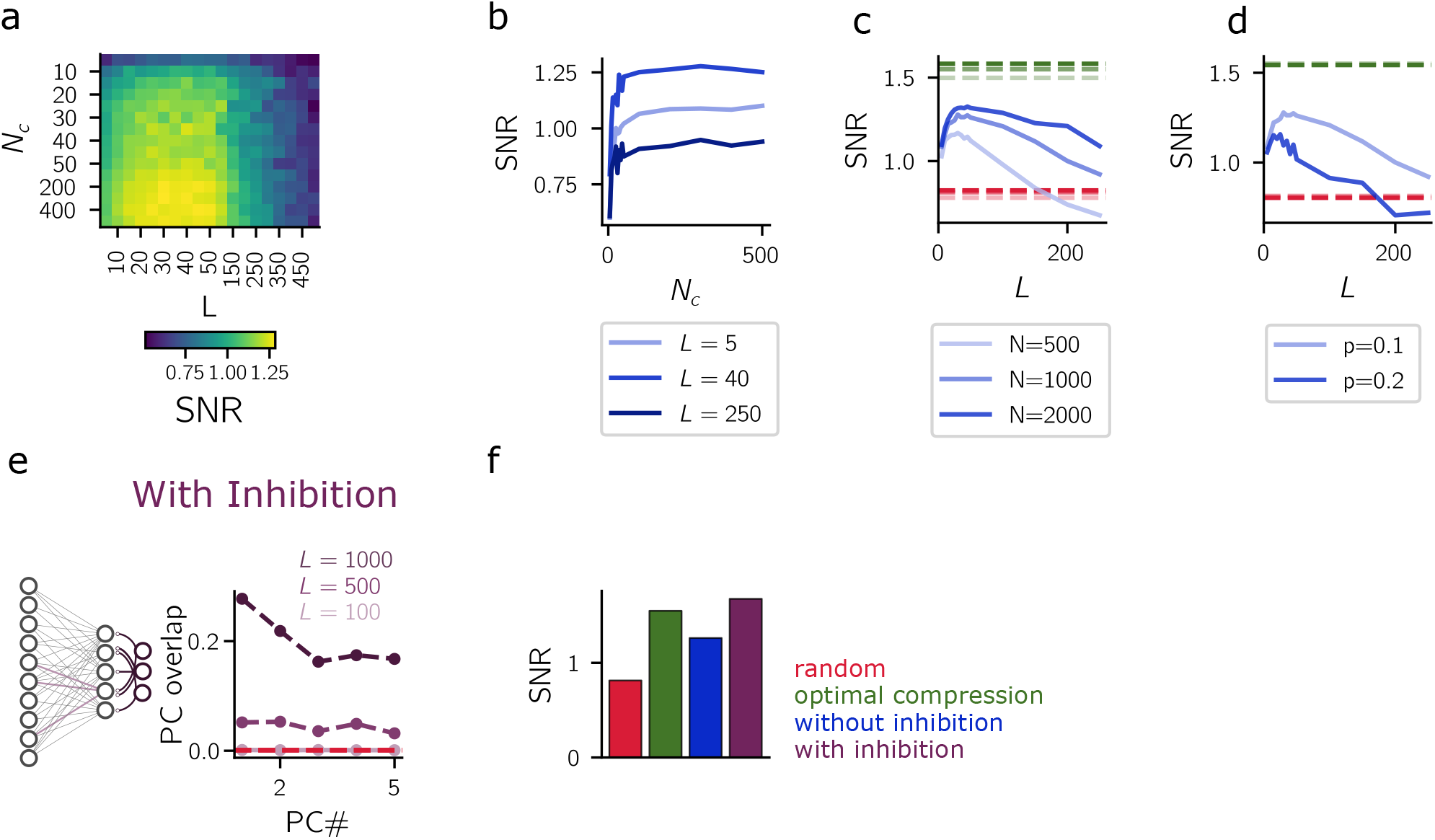
Effect of architectural paramters on Hebbian plasticity efficacy. **a, b:** Dependence of SNR on *N_c_*. The non-monotonic profile of the SNR with *L* is present for a large range of *N_c_*. **a:** heatmap and **b:** vertical slices from *L*. **c:** The SNR depends weakly on the number of input units *N*. **d:** SNR versus *L* for different values of *p*. **e:** Left: schematics of the setup in which compression weights are learned with Hebbian and anti-Hebbian learning rules in the presence of recurrent inhibition. We used the learning rule proposed in [30] (see their Eq. 18) to learn the compression weights. This learning scheme updates both the feedforward (excitatory/inhibitory) and the recurrent (inhibitory only) weights to introduce competition among compression layer units, enabling the extraction of sub-leading PCs. Right: resulting mean squared overlaps as a function of PC number as in Fig. 6a. For very large in-degree, several PCs are estimated considerably better than without recurrent inhibition. **f:** SNR at the mixed layer for different compression strategies. Biologically plausible learning achieves SNR which is statistically indistinguishable from the one of PCA compression.

**Figure S7:**
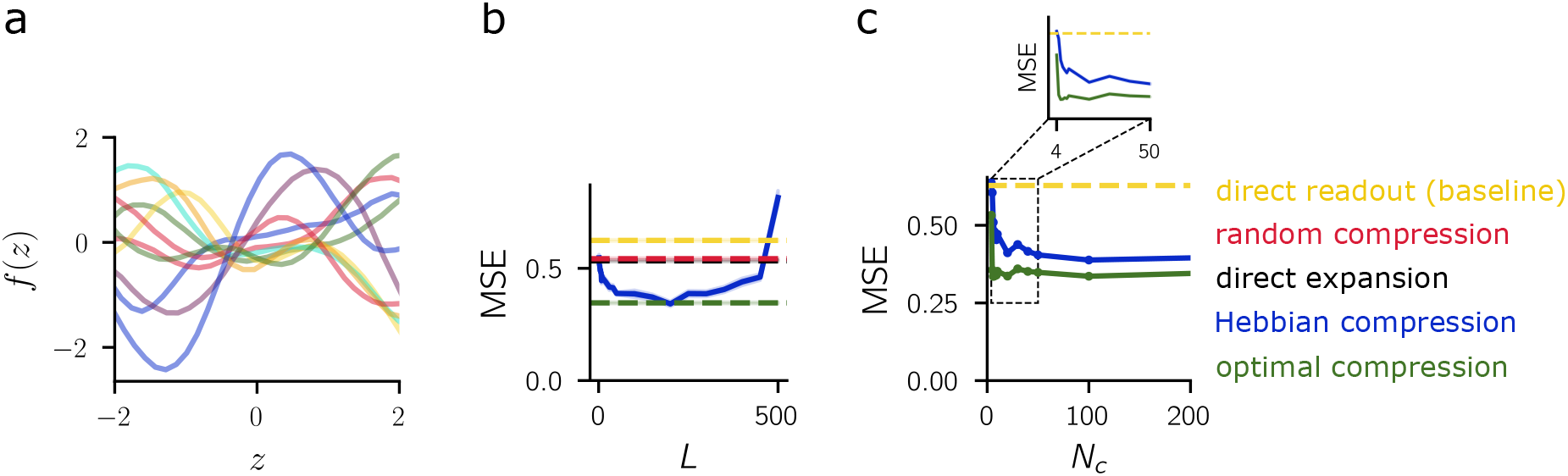
Learning random target functions. **a:** Examples of random target functions sampled from a Gaussian process prior. **b:** MSE of our model with Hebbian compression at the cortico-pontine synapses in learning a smooth Gaussian-process (GP) target, plotted against the pontine in-degree *L*. Similar to the results for classification in Fig. 6, the performance is non-monotonic with *L*. **c:** Same as **b**, but plotted against *N_c_*. The inset magnifies the small-*N_c_* region to highlight the faster decay of the MSE for optimal compression compared to Hebbian compression.

**Figure S8:**
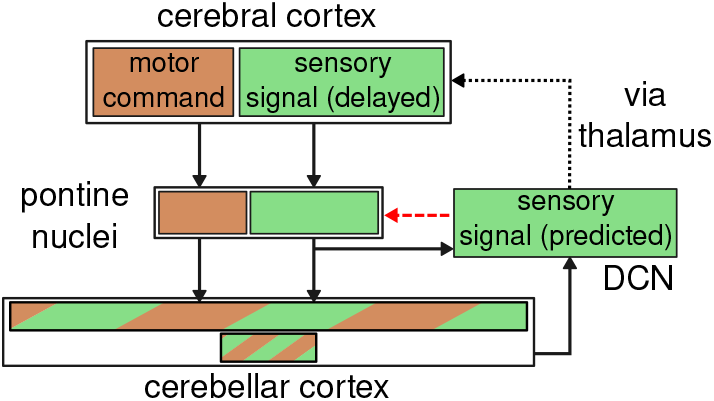
Forward model. Schematic illustrating information flow in the mammalian cortico-cerebellar pathway. We assume that the cortical representation can be segregated into motor and sensory related components. Sensory information the cortex is delayed due to sensory delays. Both representations are sent to the pontine nuclei and then mixed in the cerebellar cortex. The pontine nuclei also receive feedback (red dashed arrow) from the deep cerebellar nuclei (DCN), the output structure of the cerebellum.

